# Ligand geometry dictates cellular and *in vivo* uptake of 3D DNA nanostructures

**DOI:** 10.1101/2021.10.19.465062

**Authors:** Anjali Rajwar Gada, Shravani RS, Payal Vaswani, Vinod Morya, Amlan Barai, Shamik Sen, Sharad Gupta, Mahendra Sonawane, Dhiraj Bhatia

## Abstract

Fabrication of nanoscale DNA devices to generate 3D nano-objects with precise control of shape, size, and presentation of ligands has shown tremendous potential for therapeutic applications. The interactions between different topologies of 3D DNA nanostructures and the cell membranes are crucial for designing efficient tools for interfacing DNA devices with biological systems. The practical applications of these DNA nanocages are still limited in cellular and biological systems owing to the limited understanding of interactions of different surface topologies of DNA nanodevices with cell membranes. The correlation between the geometry of DNA nanostructures and their internalization efficiency remains elusive. We investigated the influence of the shape and size of 3D DNA nanostructure on their cellular internalization efficiency. We found that of different geometries designed, one particular geometry, i.e., the tetrahedral shape, is more favoured over other geometries for their cellular uptake in 2D and 3D cell models. This is also replicable for cellular processes like 3D cell invasion assays in 3D spheroid models and passing the epithelial barriers in *in-vivo* zebrafish model systems. Our work establishes ground rules for the rational designing of DNA nanodevices for their upcoming biological and biomedical applications.

## 1. Introduction

DNA nanotechnology utilizes DNA as a structural material to construct DNA-based nanostructures with varying designs (shapes & sizes) by harnessing fundamental properties of DNA like orthogonality in base pairing and the ability to self-assemble into higher-order structures^1–3^. Structural DNA Nanotechnology has witnessed the ground-breaking developments in recent years in terms of realizing the nano-objects of practically any shape. Recent developments in software for structure prediction can not only guide scientists to develop DNA-based objects of different shapes and sizes but also in quantitative yields and minimum off-targets. DNA nanostructures (DNs) have served as promising candidates for various biological and biomedical applications like drug delivery, biosensing, and other therapeutic applications owing to their inherent biocompatibility and low cytotoxicity^4–6^. Structural DNA Nanotechnology utilizes small strands of DNA to create various 1D, 2D, and 3D nanodevices. DNA nanostructures are gaining increasing attention in scientific research owing to their ability to be coupled to external biological targeting entities like peptides, small molecules, antibodies, etc., and their ability to encapsulate various nanoscale cargo within their internal void^7–9^. Despite such attractions, the actual applications of these devices have been limited in cellular and biological systems due to a lack of detailed investigations of such nanostructures within cellular and in vivo systems.

Cellular delivery of DNA nanostructures is gaining interest because once administered, these nanostructures must be internalized by cells in order to carry out their desired function. The study by Tuberfield and colleagues demonstrating the internalization of DNA tetrahedron in live HEK cells opened new avenues for utilizing DNA nanostructures for bioimaging and delivery of therapeutic agents into live cells^10^. The influence of geometry and size in the internalization of DNA nanostructures has been partially explored^11^. DNA origami nanostructures (DONs) with various shapes and sizes have been employed to deliver small molecules such as doxorubicin^12,13^, different proteins like nucleolin^14^, antibody^15–17^, and therapeutic nucleic acids like aptamer^18^, siRNA^19^, CpG oligonucleotides^20^ into cells. However, significantly less research has been undertaken to reveal the influence of shape, size, and cell type in the cellular uptake process. The correlation between the ligand’s topology and their internalization efficiency remains unexplored. Most of the research undertaken to investigate the design (shape & size) specific effect of DNA nanostructures has used origami-based structures that have very high molecular weight compared to 3D DNA polyheras^21,22^.

We have studied herewith the interaction of 3D DNA nanocages of different geometries and sizes with the plasma membranes of cells and their uptake pathways in cells, tissues, and animals. We have explored different geometries of DNA polyhedras like DNA Tetrahedron (TD), Icosahedron (ID), Cube, and Buckyball (BB) and compared their cellular internalization. We have also studied the correlation between geometry and cell type specific internalization in multiple cell lines derived from various body tissues. To further understand the mechanism of cellular uptake of these DNA nanostructures, we have performed inhibitor studies to identify their endocytic route. Finally, we have demonstrated the internalization of DNA nanostructures using 3D spheroid model and in the zebrafish embryos. We observe that of all the geometries explored, cell uptake is maximal for the case of tetrahedral nanocages. We also dissected the mechanism of endocytosis of these cages and found that uptake of DNA cages occurs via clathrin-mediated endocytosis. Once getting uptaken into the cells, these cages trigger processes like 3D invasion in spheroid models. This is also replicable in in-vivo model systems, where we find that tetrahedral DNA nanocages can efficiently cross the mucus barrier for their uptake into the peridermal or outermost epithelial cells in zebrafish embryos. This is for the first time, we show the demonstration of entry of DNA nanocages of different geometries and their effect of interactions with membrane, their specific endocytic uptake into the cells and their effect on cellular physiology like invasion in 3D and lastly crossing the epithelial barriers for successful uptake in model animals. Our studies will establish ground rules for future investigations involving DNA nanocages for biological and biomedical applications specifically involving their surface topologies in the areas of bioimaging, drug delivery, immune activation, differentiations, etc.

## 2. Results and discussions

### 2.1. Design, synthesis, characterization, and stability of DNA nanostructures

DNA nanostructures with different geometries were synthesized by utilizing the self-assembly property of the DNA molecule. Using previously published protocols, a panel of four structures comprising of different 3D polyhedral geometries like tetrahedron (TD)^23^, Icosahedron (ID)^24^, Cube^25^, and buckyball (BB)^26^ were designed and constructed using thermal annealing protocol in reaction buffer containing 2mM MgCl_2_ in nuclease free water. **(Figure 1a).** DNA TD was assembled using a single-step assembly approach from the equimolar ratio of four single-stranded oligonucleotides mixed in reaction buffer and annealed in the thermocycler. The reaction mixture was heated at 95°C and then gradually cooled to 4°C^27^. DNA ID utilized a modular assembly approach where different structures were synthesized separately and assembled to form a complete structure^24^. Cube and BB were assembled using three-point star motif tiles as a building block^25,26^. All the structures were labelled with cyanine-3 (Cy3) dye at their 5’end using one of the Cy3 modified ssDNA incorporated during the assembly process. DNA nanostructures were purified by gel electrophoresis and size-exclusion chromatography.

**Figure 1.**
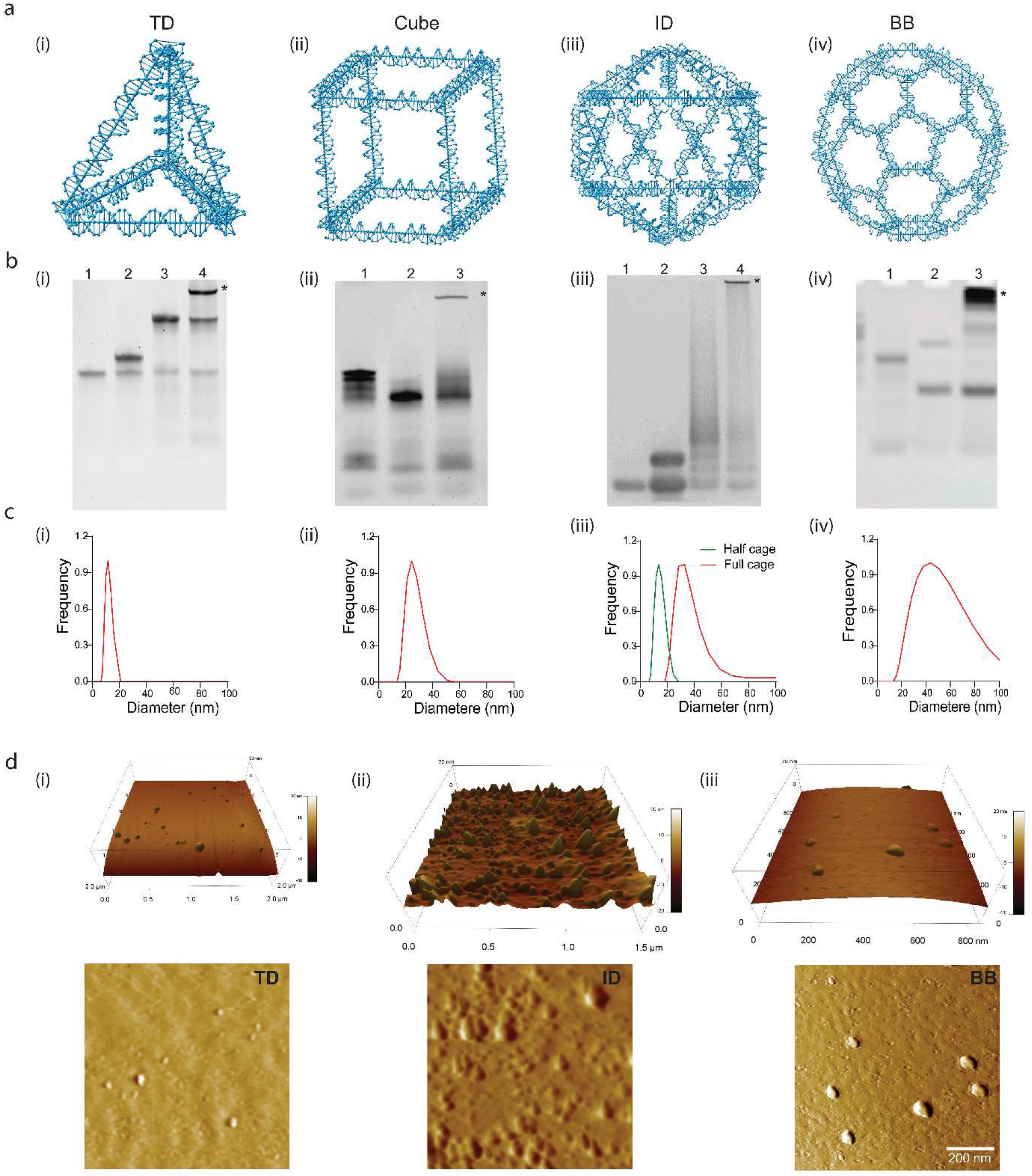
Design, Preparation, and confirmation of DNA nanostructures. **(a)** Schematic illustration DNA Tetrahedron (TD), Cube, Icosahedron (ID), and Buckyball (BB) drawn using software Nanoengineer. **(b)** Gel-based characterization showing the retardation in the mobility upon formation of ordered 3D structure. (i) Formation of TD from four strands. Lane1: T1, Lane2: T1+T2, Lane3: T1+T2+T3, Lane4: T1+T2+T3+T4 (TD) (ii) Tile-based assembly of DNA Cube. Lane1: Tile A, Lane2: Tile B, and Lane3: cube Tile (A+B). (iii) Modular assembly of ID. Lane1: ss DNA, Lane2: V5 (5-WJ), Lane3: VU5 (half modules), Lane4: Full ID (VU5 + VL5). (iv) Formation of BB from L, M, and S strands. Lane1: L strand, Lane2: L+M, Lane3: L+M+S. (*) indicates complete structure. **(c)** DLS-based characterization showing the hydrodynamic diameter of structures formed shown in figure 1b indicating the correct sizes of the objects viz., TD ~14.7nm, ID ~32.7, Cube ~24.4, and BB ~53.8nm. **(d)** Representative AFM images of (i) TD, (ii) ID, and (iii) BB showing the topology of the objects formed in figure 1b. For quantifying the sizes, 5-10 particles were analysed using Atomic J software, and the size of TD was ~8nm, ID ~28nm, and BB ~48nm. Scale Bar: 200nm.

Electrophoretic mobility shift assay confirmed the formation of the structures with the desired stoichiometry of the component oligos due to its retarding movement in the gel **(Figure 1b**). AFM and DLS were explored to test if the bands with a lower mobility in gels correspond to the objects of desired size and dimensions. The hydrodynamic size and distribution of nanostructures were detected by dynamic light scattering (DLS). DLS studies showed the hydrodynamic diameter of TDs is 14.7± 3.8nm, 32.7 ± 3.2nm for IDs, 24.4 ± 2.4nm for Cubes, and 53.8 ± 8.6nm for BBs. These values fit perfectly well with the previously reported data of these structures **(Figure 1c)**. The morphological characterization of the nanostructures was done using atomic force microscopy (AFM), which showed the characteristic shapes and sizes of the nanocages like TD, ID, and BB **(Figure 1d)**.

The stability of DNs is a very crucial parameter for utilizing these nanostructures in cellular and biological applications. DNA-based nanostructures are highly susceptible to nuclease degradation outside and inside the cells and can give deceptive fluorescence signals. To test the stability of DNs to the cellular environment, we incubated all DNs with 10% FBS, the component carrying nucleases. We found that all the DNs were stable for up to 1 hour. (**Figure S1** in supplementary information).

### 2.2. Concentration-dependent uptake of DNA nanostructures in mammalian cells

Cellular uptake of DNA constructs relies on transfection agents aid that helps transport DNA across the negatively charged plasma membrane. However, DNA-based nanostructures are known to traverse the membrane of mammalian cells without any aid of transfection agents. Intuitively, we have studied the cellular uptake of well-defined 3D DNA nanostructures (DNs) with different topologies in the MDA-MB-231 breast cancer cell line. A concentration-dependent study (5 nM to 500 nM) was performed to identify the optimum concentration for further cellular studies. Cells were seeded 24h before the experiment in DMEM media supplemented with 10% FBS. We studied the uptake of single-Cy3 labelled DNs and Alexa488 labelled transferrin (Tf-A488) for 20 min at 37°C. Tf-A488 is an endocytic marker of clathrin-mediated endocytosis that served as an internal control. The concentration of Tf-A488 used was 5 μg/ml and kept constant in all the experiments. The non-treated cells were considered as a negative control for validating the internalization efficiency of DNs.

The internalization of nanostructure was characterized both qualitatively and quantitatively using laser scanning confocal microscopy. The internalization of DNs was tracked using Cy3 conjugated DNA, which was incorporated during the assembly reaction. The concentration of Cy3-DNA was constant in TD and ID, whereas the number of copies of Cy3 DNA in cube and BB was more, which was normalized by dividing the total intensity by the number of fluorophores. The non-treated cells showed negligible signal in the red channel, validating that the fluorescence in the treated cells corresponds to the internalized DNs. For TD, the signal can be detected at concentrations as low as 5 nM **(Figure 2a, b),** whereas for cube and BB the signal was visible only after 25 nM and 10 nM, respectively (Supplementary information, **Figure S2, and S3**). However, ID’s signal was significantly less than the other DNs **(Figure 2c,d)**. The signal of TD increases linearly with the concentration, whereas for Cube and BB, the signal increases till 250 nM and then decreases which may be due to either the aggregation of DNs at higher concentrations or self-inhibition on the membrane due to crowding. The images were processed and quantified using ImageJ software. We further conducted a time-dependent study (0, 5,10, 15, 30, and 60 min) in parallel to study the cellular uptake of DNA TD over time. Initially, the intensity of TD from cells increased with time up to 15 mins, after which it started to decrease indicating the recycling of the internalized TD from the cells **(Figure 2e,f)**.

**Figure 2.**
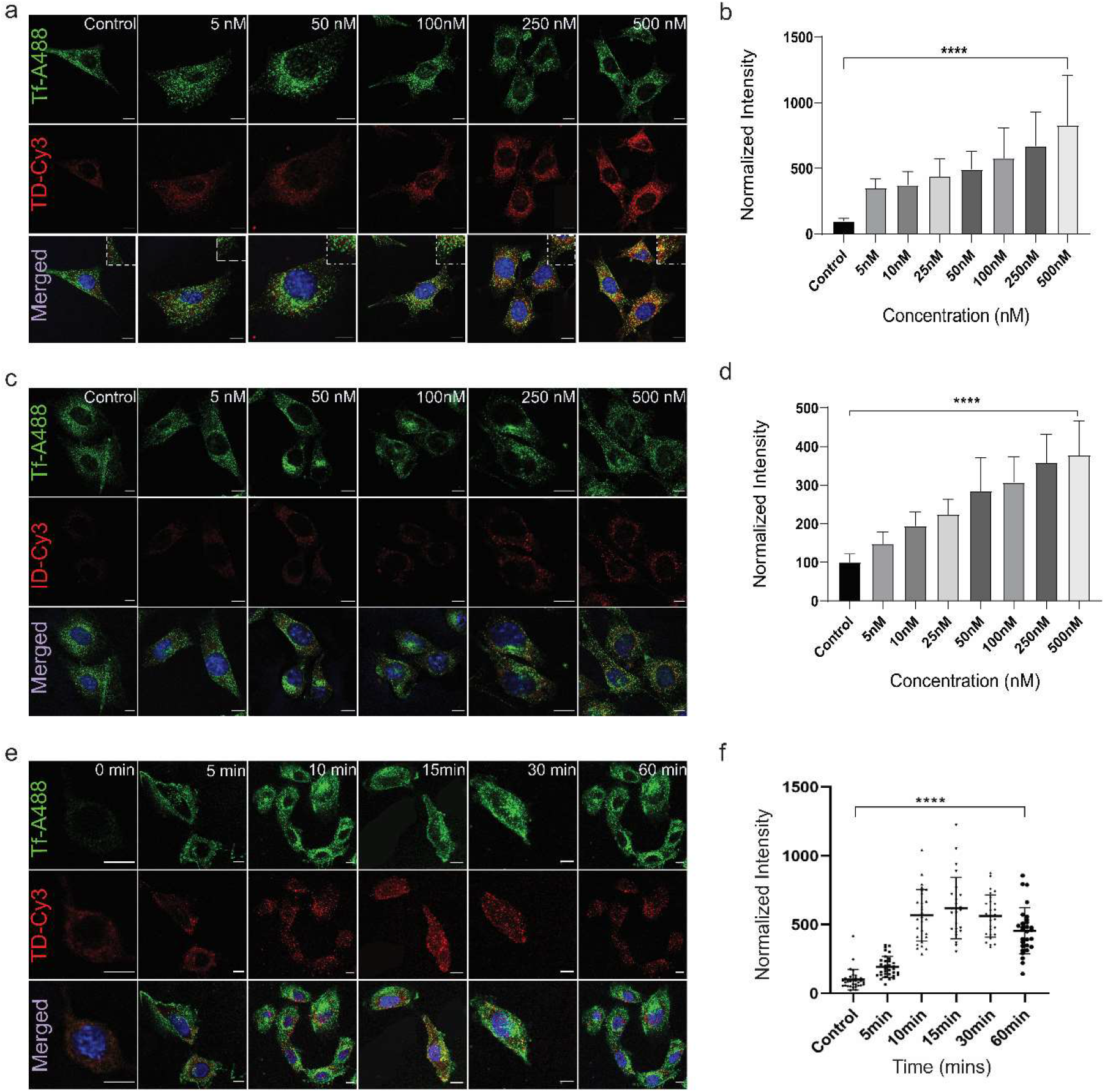
Cellular uptake of DNA nanostructures. **(a,c)** Confocal images of MDA-MB 231 cells treated with DNA TD-Cy3 and ID-Cy3 in different concentrations ranging from 5 nM to 500 nM. Green channel shows the uptake of TF-A488 (internal control), Red channel represents TD-Cy3 and ID-Cy3 uptake and the bottom panel represents the merged images of all the channels with nuclei stained with Hoechst dye. **(b,d)** Quantification of TD-Cy3 and ID-Cy3 uptake in MDA-MB231 cells from panel (a & c) respectively. **(e)** Time dependent study of DNA TD from 0 min to 60 min. Green channel represents TF-A488, Red channel: TD-Cy3 at different time points. **(f)** Quantification of TD-Cy3 uptake from cells in panel (e). Error bars in all quantification indicate standard errors. The normalized intensity was calculated from 40 cells (ordinary one-way ANOVA, P value < 0.0001****). Scale Bar: 10μm.

### 2.3. Cellular uptake of DNs is size and cell type dependent

How different DNs bearing different topologies interact with complex cellular environments of different cell types can be utilized to provide a unified piece of information about the cell-specific preference of DNA nanostructures. We studied the uptake of DNs in a panel of cell lines originating from different sources and body tissues including SH-SY5Y, HeLa, SUM159, KB3, and RPe1. We found 150 nM is the optimum concentration to study the cellular uptake from the previous experiment and was used for further experiments. DNs of different geometries were uptaken with varying efficiencies in SUM159 cells **(Figure 3a, b)** and other cell lines as well (Supplementary information, **Figure S4, S5, S6 and S7)**. We found that DNA TD’s uptake was higher than other DNs in all the cell lines. We further compared the internalization of TD in different cell lines to investigate if there is any tissue specific preference in the internalization of DNA TD. We found that the internalization of TD was more in the cancerous cell lines compared to the non-cancerous cell lines. Among the cancerous cell lines the internalization was maximum in breast (SUM159), kidney (HEK293T) and liver (WRL) cell lines **(Figure S8)**.

**Figure 3.**
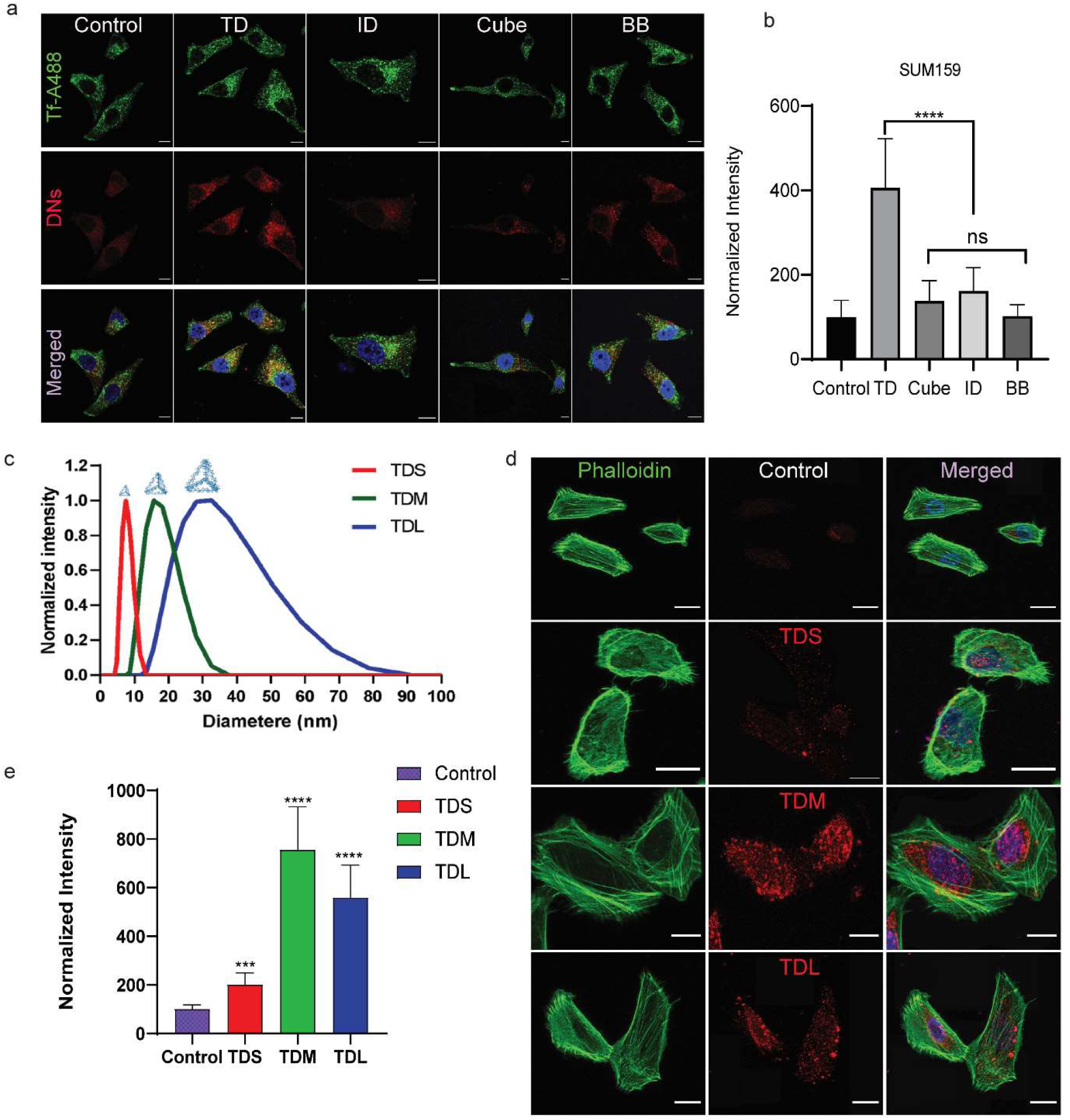
Effect of cell type and size in the uptake of DNA Nanostructures. **(a)** Cellular uptake of TD, ID, Cube, BB in SUM 159 cells for 20 mins at 37°C. Confocal images of cells showing Green channel represents the uptake of Tf-A488, Red Channel represents Cy3 labelled nanostructure and bottom panel represents the merged image with nuclei stained with Hoechst. **(b)** Quantification of DNA nanostructure uptake in SUM159 cells from panel (a). Error bars in the quantification indicate standard errors. The normalized intensity was calculated from 40 cells (ordinary one-way ANOVA, P value < 0.0001****). **(c)** DLS based characterization of different sizes of DNA TD. The hydrodynamic diameter of small TD (TDS) ~11.7nm, medium TD (TDM) ~14.3nm and large TD (TDL) is ~32.7nm. **(d)** Confocal images of SUM159 cells show DNA TD’s size-dependent uptake at 150nM concentration for 20mins at 37°C. Green channel represents actin cytoskeleton stained with Phalloidin-A488, red channel represents the different size of DNA TD, and the bottom panel represents the merged image with nuclei stained with Hoechst. **(e)** Quantification of the DNA TD internalization from cells in panel (d). TDM and TDL showed more internalization compared to TDS. Error bars indicate the mean of the two experiments with the associated s.d. (two-tailed unpaired t-test. **** p < 0.0001, * n = 2). Scale Bar: 10μm.

Since DNA TDs were getting internalized in all the cell types with maximum efficiency, we further investigated if the size of TD has any role to play in the internalization of DNA TDs. TDs of three different sizes were synthesized using thermal annealing protocol as described previously. The hydrodynamic size of TD was analysed by dynamic light scattering and found to be 11.7 nm for small TDs (referred to as TDS), 14.3 nm for medium TDs (referred to as TDM), and 32.7 nm for large TDs (referred to as TDL) **(Figure 3c)**. We did the cellular uptake studies in the SUM159 cell line at a 150 nM concentration of TD. We hypothesized that TDS will be internalized more owing to its smaller size but in contrast, we found that TDM followed by TDL has shown maximum internalization **(Figure 3d, e)**. This explains that the size of the ligand indeed has a vital role to play in the internalization of DNA nanostructures.

### 2.4. DNA TDs get endocytosed by Clathrin Mediated Endocytosis

Efficient cellular uptake of the cargo is the primary requirement for delivering molecules of interest. We optimized the uptake of different DNs and concluded that of the tested geometries, TDs are most efficiently internalized in all the cell types. We further wanted to delineate the cellular pathways involved in the internalization of DNA TDs. The uptake of small ligands into cells takes place mainly via active processes like endocytosis. There are two major pathways for endocytosis broadly classified into clathrin-mediated endocytosis (CME) and clathrin-independent endocytosis (CIE) involving the clathrin machinery. Alternatively, less efficient modes such as phagocytosis are involved in the uptake of large sized structures like bacteria. However, these are not very efficient and very slow in action. Since DNA TDs are ligands of size 15nm, we ruled out the phagocytosis and used inhibitors to block the CME and CIE pathways. Pitstop-2 is a well-studied inhibitor of the CME pathway, which interferes with the terminal domain of clathrin involved in coated pit dynamics, thereby inhibiting clathrin pit scission from the membrane and blocking vesicle transport to the endosome^28^. Lactose has been shown to compete with lectins like galectin-3 which binds the glycosylated receptors thereby blocking the formation and uptake of clathrin independent carriers^29^.

Cells were pre-treated with two different concentrations of Pitstop-2 (20μM & 40μM) to block CME process for 15mins at 37°C specifically. Control cells and treated cells were probed to check the uptake of TD-Cy3, Tf-A488, and Gal3-Cy5 after 20 min of incubation at 37°C. For pitstop-2 treatment, two different concentrations were tested, and the pathway got inhibited at 20 μM; however, the cells got stressed at 40 μM. Transferrin is a well-established marker for the CME pathway and thereby was used as a positive control. Galectin-3—known to enter the cells via the CIE pathway—was used as a negative control for Pitstop-2. The CME pathway got inhibited upon treatment with 20 μM of Pitstop-2, as confirmed by the quantitative reduction in the signal of Tf-A488, whereas the uptake of Gal3-Cy5 was not significantly affected. The signal from TD was significantly reduced, suggesting the internalization of TD by the CME pathway **(Figure 4a, b).**

**Figure 4.**
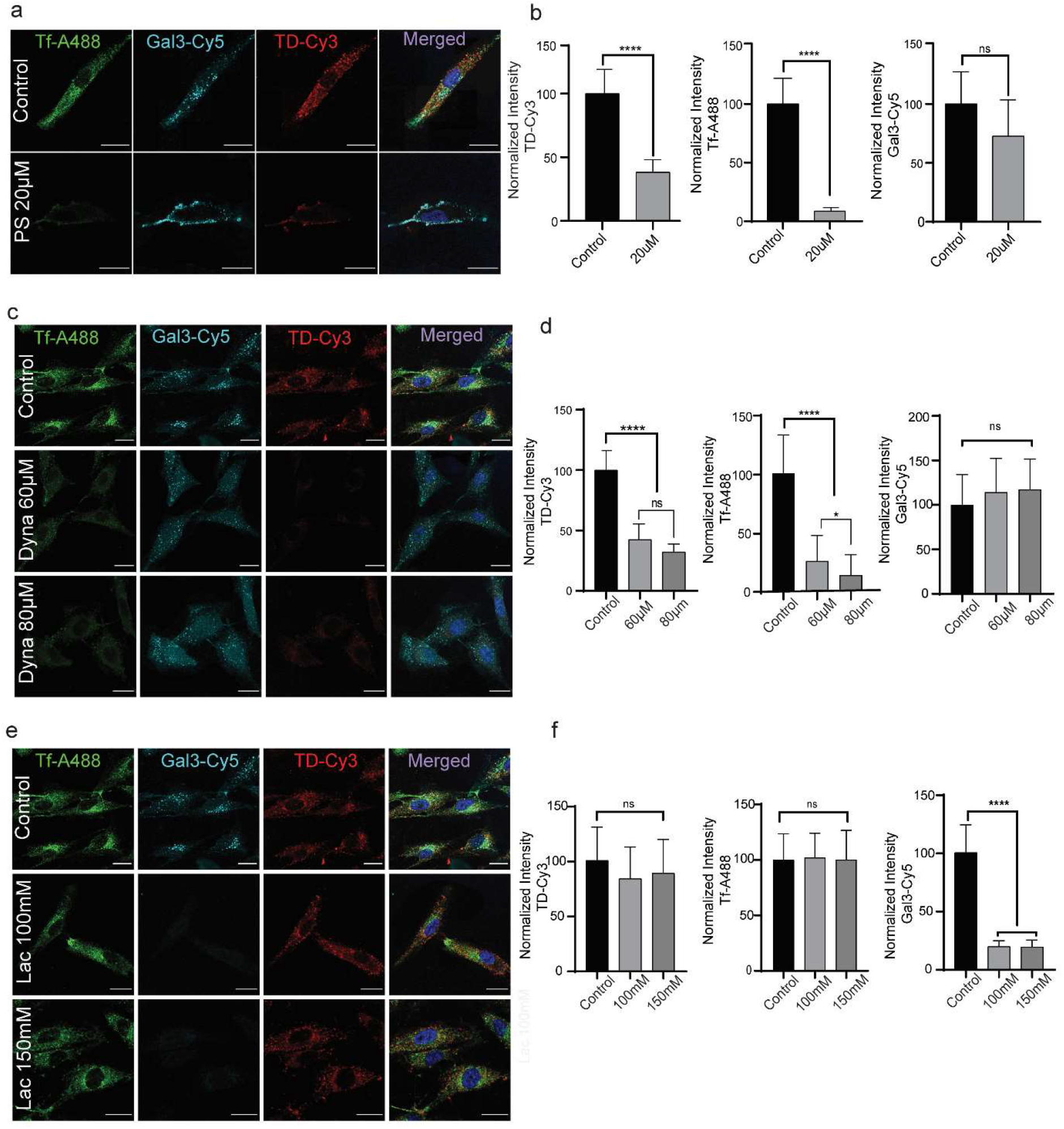
DNA TDs get endocytosed by clathrin mediated endocytosis in mammalian cells. **(a)** Confocal images of Rpe1 cells showing the uptake of Tf-A488 in green channel, Galectin3-Cy5 in Cyan channel, and TD-Cy3 in red channel for 20 min at 37°C in the presence or absence of 20 μM Pitstop-2. **(b)** Quantification of inhibition of CME pathway by Pitstop-2 for Tf-A488, TD-Cy3 and Gal3-Cy5 from cells in panel (a). Significant inhibition of TD and Tf but not significant inhibition of Gal3. **(c)** Blocking the dynamin-dependent pathway by inhibiting the dynamin via dynasore inhibitor at 40 μM and 60 μM concentrations. The cellular uptake of TF-A488, Gal3-Cy5 and TD-Cy3 was monitored with and without the treatment of dynasore for 20 min at 37°C. **(d)** Quantification of inhibition by measuring the fluorescence intensity of TF-A488, TD-Cy3 and Gal3-Cy5 with and without treatment for dynasore from cells in panel (c). The uptake of Td and Tf were significantly inhibited at both the concentration whereas the uptake of Gal-3 was not very significantly affected. (e) Uptake of Tf-A488, Gal3-Cy5 and TD-Cy3 with inhibition of CIE pathway by Lactose. (f) Quantification of inhibition of CIE pathway for Tf-A488, Gal3-Cy5 and TD-Cy3. Significant inhibition of Gal3-cy5 but no significant inhibition of TD-Cy3 and Tf-A488. For all quantifications, error bars indicate the mean with associated s.d. (two-tailed unpaired t-test. **** p < 0.0001, * p<0.05, ns: non-significant). Scale Bar: 10μm

We used another inhibitor, ‘Dynasore’ to block the CME pathway to further confirm the above results. Dynasore inhibits dynamin that acts as a molecular scissor and regulates the dynamics of clathrin-coated vesicles^30^. Dynaosre inhibits the GTPase activity of dynamin which is involved in the pinching of the vesicles. The CME pathway was blocked by pre-treating the cells with 60 μM and 80 μM concentrations of Dynasore for 30 min. Again, TD uptake was significantly affected along with Tf-A488, suggesting CM-dependent endocytosis of TDs **(Figure 4 c,d)**. However, the internalization of Gal-3 was not severely affected. In contrast, the uptake of TDs remains unaffected mainly in cells pre-treated with lactose, a known competitive inhibitor of galectins that blocks the CIE pathway^31^ **(Figure4 e,f)**. Taken together, these findings support a CME-dependent mechanism of TD internalization. We additionally checked the internalization route of TD in MDA-MB231 cell lines and observed the similar trend results i.e., upon treatment with pitstop-2 and dynasore the internalization of TD was significantly reduced. In contrast, upon lactose treatment the internalization of TD was unaffected inferring the internalization of TD via CME process **(Figure S9, S10)**. Previous reports suggested the internalization of many DNs via the caveolin pathway. We employed methyl beta-cyclodextrin (m*β*CD) inhibitor to selectively block the caveolin pathway^32^. We found that the internalization of TDs was not significantly disturbed upon treatment with m*β*CD whereas the uptake of CtxB, a potential cargo for caveolin pathway was selectively inhibited^33^(Supplementary information, **Figure S11**). However, caveolin mediated uptake is present mainly in muscle cells and its expression is minimum in other body cells whereas clathrin mediated and independent pathways are ubiquitous in all body cell types.

### 2.5. DNA TDs stimulate 3D cell invasion in spheroid model and in vivo uptake in zebrafish model system

3D cell culture mimics cell’s physiological environment and thus provides an improved platform to study cellular spreading, migration, invasion as well as differentiation. In 3D cell culture, cells clump together to form stable cell-to-cell contacts, thereby maximizing the communication and signalling between the cells. MDA-MB-231, extremely aggressive triple-negative human breast cancer cells, were used to prepare 3D spheroids using the hanging drop method. It is a non-scaffold-based method in which cell suspensions are placed dropwise on the lid of the petridish^34^. The cells aggregated due to gravity and form a compact spheroid at the base of the drop after 36 hours of culture at 37°C. These spheroids can then be transferred to the collagen matrix to invade the 3D system. This invasion potential of the cells can be either stimulated or inhibited by using specific activators or inhibitors. We wanted to check if DNA nanostructures had any effect on the invasion potential of these spheroids. Spheroid invasion assay provides an attractive *ex vivo* platform to study the invasion potential of 3D DNA-based nanodevices. Cell invasion potential can be quantified by measuring the dispersion of cells in different directions or by calculating the migration distance covered by cells from the intact spheroid core. Following the characterization of uptake of different DNs in the 2D cell culture system, we then investigated the uptake of DNs in the 3D spheroid models. To evaluate the invasion potential of DNs with different topologies, we incubated the spheroids with DNs for 24 h at 37°C. Untreated spheroids were considered as control that showed minimal invasion potential as the migration distance was negligible, and hardly any cells invaded out from the spheroid core into the collagen matrix **(Figure 5a).** The zoom-in image of the section of spheroids also demonstrated successful endocytosis of the DNA TDs into migrating cells, thereby stimulating their invasion (**Figure 5b**). Spheroids treated with DNs showed visible cell invasion in the matrix, the migration distance was more, and many cells migrated in different directions away from the spheroid core. Among all, TD showed the maximum invasion, followed by ID, BB and Cube **(Figure 5c)**.

**Figure 5.**
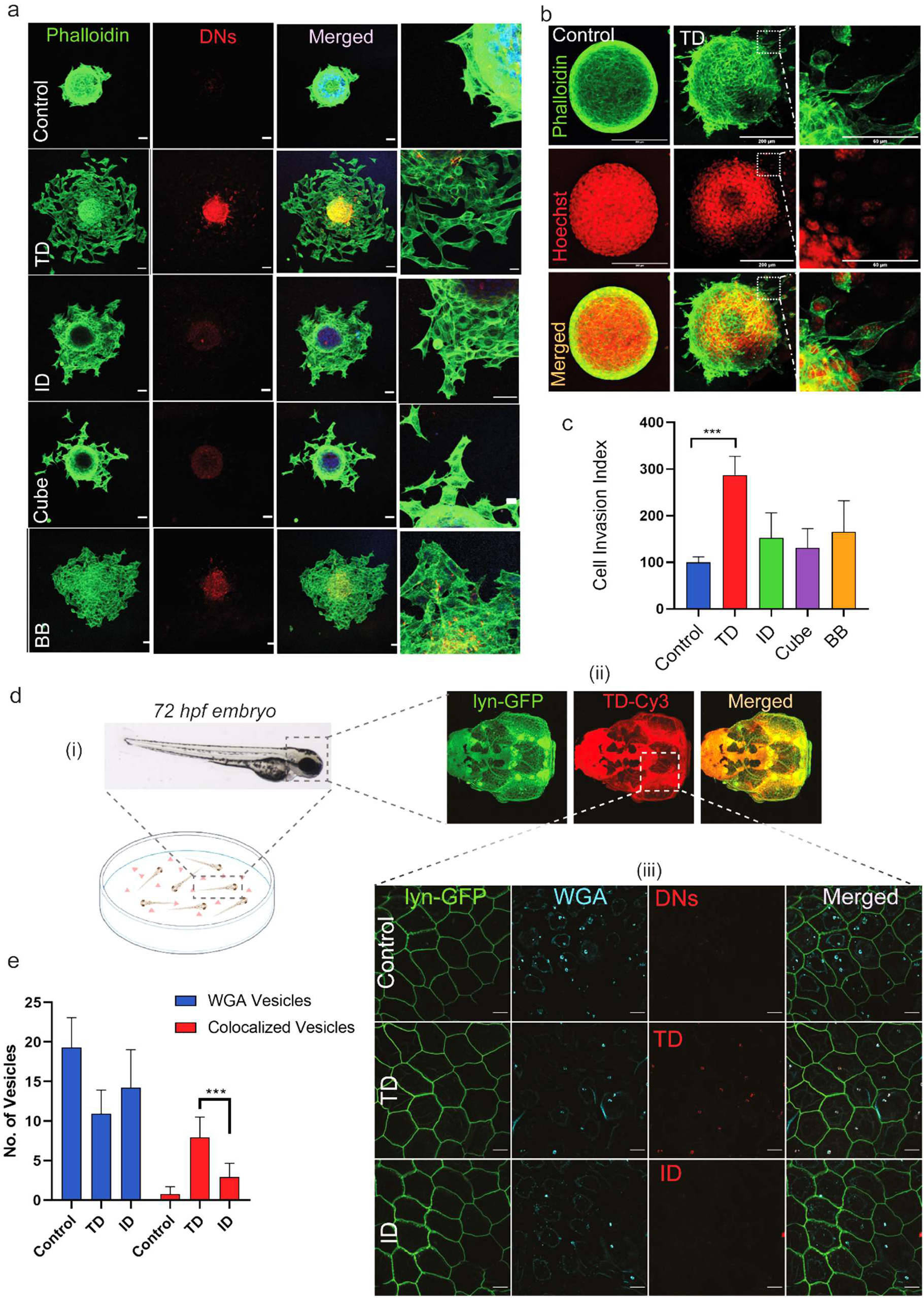
DNA TDs stimulate 3D cell invasion in spheroid model and in vivo uptake in zebrafish model system. (a) Confocal images of DNs uptake by MDA-MB231 spheroids after 24 hours of incubation at 37°C. DNs have shown invasion in 3D spheroids as cells showed migration from the spheroid core. The green colour represents Phalloidin staining of actin filaments, red colour represents Cy3 labelled DNs. Scale Bar: 100μm. (b) Magnified images of 3D spheroids. Non-treated spheroids were considered as control. TD has shown maximum invasion followed by BB. Scale Bar: 200μm (c) Quantification of cellular invasion from spheroids shown in panel (a). Cell invasion index was calculated by measuring the migration distance by cells from the spheroid core. For each nanostructure, a minimum of 8 spheroids were quantified. Error bars indicate Standard error (ordinary one-way ANOVA, p-value < 0.0001****). (d) In-vivo uptake of DNs in the zebrafish embryo. (i) Schematic of experimental design. 72 hpf zebrafish embryos were soaked in media containing 300nM of DN and incubated for four hours. (ii) Confocal images of the dissected head of 72hpf embryo showing the internalization of TD shown in red colour. Green colour represents membrane-tethered lynEGFP. (iii) Confocal images of the peridermal cells of zebrafish embryo showing the internalization of DNs. Scale Bar: 10μm (e) Quantification of DNs internalized into the peridermal cells, calculated by counting the number of DNs vesicles colocalized with WGA vesicles which is known to mark endocytosis in general. Error bars indicate the mean with associated s.d. (two-tailed unpaired t-test. *** p < 0.001). For each condition n=6-8 embryos were quantified and (N)=2.

**Figure 6.**
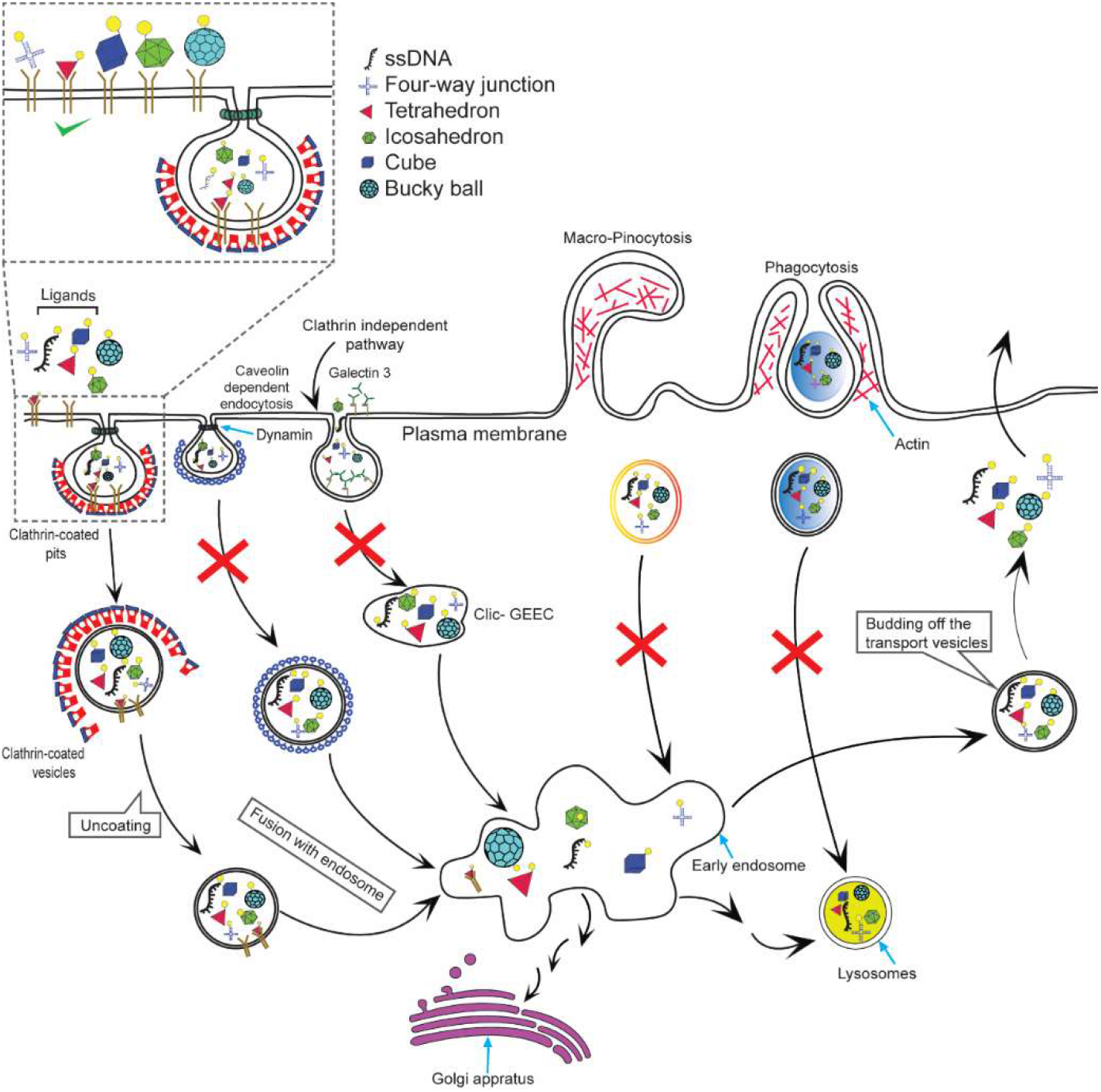
Schematic illustration of the internalization pathway potentially followed by Tetrahedral DNA in cells. Multiple pathways co-exist at plasma membrane for internalization of various cargos including nutirents, growth factors, toxins and even fluid. We successfully marked two different endocytic pathways viz clathrin dependent and independent pathways and we found that DNA nanocages of different geometries mostly adopt the clathrin mediated endocytosis. Further, the uptake of these nanocages is strongly correlated with the object’s geometry and found that only tetrahedral geometry is more preferred for uptake in cells and in vivo.

We further asked if the geometry of DNs has a similar impact on their uptake in the in-vivo systems as was observed in cells. We tested this by performing an in vivo uptake assay of TD and ID in zebrafish embryos, as these DNs showed the maximum invasion potential in the previous experiment. The in vivo uptake of DNs was performed in zebrafish embryos obtained from a transgenic line *Tg(cldb::lyn-EGFP)* in which the expression of lynEGFP is driven by *claudin* promoter marking the membrane of peridermal cells. To check the entry of nanocages, 72 hpf (hours post fertilization) embryos were soaked in a solution containing 300 nM of DNs for 4h **(Figure 5d (i)),** post which the embryos were washed, fixed, and imaged using confocal microscopy. Wheat germ agglutinin (WGA) was used as an internal control to show the potential of these DNs to overcome the mucus barrier and their successful uptake into the zebrafish peridermal cells. We observed that the internalization of TD was more compared to the ID and control. (**Figure 5d (iii)**). Further quantification of the number of vesicles labelled by control, TD, and ID and colocalized with the WGA vesicles, confirmed this trend **(Figure 5e)**.

### Conclusions

The practical applications of DNA nanodevices in biological systems are minimal and that too very coarsely characterized. This is primarily due to the lack of in-depth understanding of interactions of DNA nanodevices with biological membranes and then applying the knowledge of these interactions to develop Smart Nanoparticles for biological and biomedical applications.

Herein we present the basic principles showing how the geometry and size of the DNA based nanoparticles play crucial roles in interactions of the DNs with membranes leading to their very specific uptake into mammalian cells. Most of the endocytic pathways like clathrin-mediated endocytosis trigger signalling pathways like MAPK, SRCK, etc which trigger the range of activities leading to cell migration in 2D and 3D^35,36^. It is through this active endocytosis of DNs into the cells that trigger 3D invasion of cells from spheroids into the collagen matrix. Not only do these nanoparticles activate these physiological pathways in a geometry specific manner, but they also possess the capacity to enter epithelial cells in in-vivo systems like zebrafish embryos. Taken together, our results have established the strict influence of ligand geometries on their interactions with biological systems. These results will play crucial roles in the future development of DNA based smart nanoparticles with specific and targeted capacities to probe and program biological systems in multiple avenues like bioimaging, targeted delivery, activation of immune cells, or as scaffolds for tissue engineering, to name a few.

## 3. Materials and Methods

### 3.1. Materials

Dulbecco’s Modified Eagle Medium (DMEM) was purchased from Lonza. DMEM-F12, HAMS-F12, Fetal bovine serum (FBS), PenStrap, Trypsin- EDTA (0.25%), collagen were purchased from Gibco and Phosphate buffer saline (PBS) was purchased from HyClone. The sequences used for DNA nanostructure synthesis (Table 1–5), 6X loading dye, 50 bp DNA ladder, mowiol, transferrin-A488, Cy5, Hoechst, Pitstop-2, Dynasore and Lactose were ordered from Sigma Aldrich. Nuclease free water, ammonium persulfate, ethidium bromide, TEMED, paraformaldehyde and adherent cell culture dishes were purchased from Himedia. Tris-Acetate EDTA (TAE), Acrylamide/bisacrylamide sol 30% were purchased from GeNei. Magnesium chloride was ordered from SRL, India. Galectin-3 was provided as a gift from Johannes team at Institut Curie, Paris.

### 3.2. Synthesis of DNA nanostructures

The synthesis of TD^27^ and BB^26^ was done by one pot synthesis. The primers were reconstituted in nuclease free water to 100 μM stock and diluted to 10 μM working concentration. DNA TD was synthesized using four oligonucleotides mixed in equimolar ratios (T1:T2: T3:T4 - 1:1:1:1) (Table1) containing 2mM MgCl_2_. BB was synthesised using L (long) strand, M (medium) and S (small) strands. For BB, the L:M:S ratio was 1:3:3 with 2mM MgCl_2_. The reaction was carried out in a thermocycler and the reaction mixture was heated to 95°C and then gradually cooled to 4°C. The reaction cycle has step 5°C decrease with a interval of 15min at every step.The final concentration for TD, and BB was, 1 μM and, 1.1 μM respectively.

The synthesis of ID and cube was done through modular assembly. For ID, three 5WJ are formed in the first step (V5, U5 and L5) with equimolar ratio of the primers (Table 2) and 2 mM MgCl_2_ is added. In the second step, one part of V5 is combined with 5 parts of U_5_ and L_5_, each forming two half icosahedrons (VU_5_ and VL_5_). This reaction is also catalysed by MgCl_2_. The last step is 1:1 assembly of VU_5_ and VL_5_. The first step is performed on 95°C to 4°C, similar to one pot assembly. The second and third step use the temperature 45°C decreasing to 4°C with 5°C decrease in each step for 30 minutes. For cube, the first step is creating Tile A (L:SA:MA – 1:3:3) and Tile B (L:SB: MB – 1:3:3) with 2mM MgCl_2_. The annealing conditions are like one pot reaction. The second step is 1:1 mixture of Tile A and Tile B and the reaction condition is similar to second and third step of ID assembly. The final concentration of ID and cube were 1.6 and 1.1 μM respectively. All the structures were stored at 4°C until further use.

### 3.3. Characterization of DNA Nanostructures

#### 3.3.1. Electrophoretic mobility shift assay (EMSA)

The size-based characterization was done using three methods. Electrophoretic mobility shift assay (EMSA) was performed using Native- PAGE. 5% polyacrylamide gel was prepared to study the higher order structure formation. The sample contained 5 μL of DNA nanostructure sample with 3 μL of loading buffer and 1.5 μL of 6X loading dye. The gel was run on 80 V for 80 minutes. The gel was stained with EtBr stain and visualized using Gel Documentation system (Biorad ChemiDoc^TM^ MP Imaging System).

#### 3.3.2. Dynamic Light Scattering (DLS)

The size-based characterization of DNs was done to measure the hydrodynamic size of the formed structures using Dynamic Light Scattering (DLS). The sample was diluted in a 1:20 ratio, and 50 μL of samples was used to analyze the hydrodynamic radius using the Malvern analytical Zetasizer Nano ZS instrument. The readings were taken in triplicates, with 13 readings in one run.

#### 3.3.3. Atomic Force Microscopy (AFM)

The morphology-based characterization was performed using Atomic Force Microscopy. A freshly cleaved mica was treated with 1 mM NiCl2 solution for 15 minutes and then washed 2 times with ultrapure water. 20 - 30 μl droplets DNA nanostructure samples (150nM) were then overlayed on top and was incubated for 15 minutes. The excess sample was then drained and washed 2 times with ultrapure water and was dried and imaged using MPF-3D BIO AFM (Asylum Research, Oxford Instruments) in tapping mode in air. A sharp silicon cantilever AC160TS (Olympus) was used for AFM imaging.

### 3.4. *In Vitro* studies

#### 3.4.1. Cell Culture

HeLa (Cervical Cancer), MDA-MB-231 (Breast Cancer), Rpe1 (Retinal pigment epithelial cells), HEK293T (Human Embryonic Kidney cells), KB3 (Oral Cancer) cells were maintained in DMEM, SUM-159-A (Breast Cancer) cells were maintained in HAMS-F12 media and SHSY5Y (Human Neuroblastoma) were maintained in DMEM-F12. All the media were supplemented with 10% foetal bovine serum and 1% penicillium/streptomycin. The cells were maintained at 37°C with 5% CO_2_ in a humidified incubator.

#### 3.4.2. Cellular Uptake Assay

For cellular uptake experiments, all the cells were maintained in their respective media described previously. Cells were seeded at a density of 0.1*10^6^ cells/well on 18mm glass coverslips in 12 well plates 24hrs before the experiment. Before experiments, cells were checked on the microscope to visualize their proper attachment and spreading. Cells were washed twice with 1x PBS and then incubated with serum-free media containing different concentrations of DNs and 5ug/ml of Tf-A488 for concentration-dependent and cell-type specific experiments. The cells were incubated at 37°C for 20 minutes and then washed twice with acid buffer (pH-2.5) followed by three time washing with 1x PBS to remove the excess or surface bound DNs. The cells were fixed using 4% paraformaldehyde for 15mins at 37°C. The cells were again washed thrice with 1x PBS and mounted on the glass slide using Mowiol containing Hoechst to mark the nucleus.

#### 3.4.3. Inhibitor Studies

The endocytosis inhibitor study was performed to understand the pathway by which nanostructures are being up taken. Rpe1 cells were seeded into 12-well plate on coverslips and grown till 80-90% confluency. The cells were washed with PBS and pre-incubated with Pitstop (20 μM, 40 μM) or Dynasore (60 μM, 80 μM, 100 μM) or Lactose (100 mM, 150 mM, 200 mM) in serum free DMEM for 15 minutes at 37°C. The media was decanted, and the cells were treated with the same concentration of inhibitors with transferrin-A488 (5 μg/mL), galectin3-Cy5 (5 μg/mL) and TD (150 nM) in serum free DMEM. The cells were incubated at 37°C for 15 minutes. The cells were then washed with PBS and fixed with 4% PFA at 37°C for 15 minutes. They were washed with PBS and mounted with Mowiol and Hoechst to mark the nucleus.

#### 3.4.4. 3D spheroid invasion assay

MDA-MB231 cells were used to prepare 3D spheroids using *hanging-drop method.* The cells were trypsinized and appropriate volume of complete media was added. 2.5 × 10^4^ cells were taken in total 50 μl of complete media and seeded in form of droplets on lid of petri dish. The base of petri dish was filled with 30 mL of PBS to provide humidity for spheroid growth. The cells were incubated at 37°C for 36 hours. The cells clumped together and aggregated due to gravity to form spheroids. The spheroid formation was confirmed by observing under bright field optical microscope. The spheroids were transferred to 12 well plate using 3:1 collagen to media proportion containing collagen covered coverslips and it was incubated at 37°C for 1 hour. The spheroids were then incubated with different nanostructures at 150 nM in serum free media for 24 hours at 37°C. The spheroids were fixed using 4% PFA for 15 mins at 37°C. For Phalloidin staining spheroids were first permeabilized using 0.1% TritonX and incubated for 10mins at room temperature. After that 0.1xTritonX+Phalloidin solution (1:1000 dilution) was added and incubated at 37°C for 30 mins. Spheroids were washed gently twice with 1x PBS and mounted using Mowiol+Hoechst.

#### 3.4.5. Confocal Microscopy

Fixed cells were imaged using Confocal Scanning Laser Microscope (Leica TCS SP8). The 2D cell culture assay slides were imaged using 63x oil immersion objective, while the 3D cell culture (spheroids) assay slides were imaged using 10x and 20x objective.The pinhole was kept 1 Airy unit. Different fluorophores were excited using different lasers i.e. for Hoechst: 405nm, Tf-A488: 488nm, TD-Cy3: 561nm and Gal3-Cy5 with 633nm. The image analysis was done using Fiji ImageJ software.The background from each image was subtracted using Fiji ImageJ software (NIH) and whole cell intensity was quantified using maximum intensity projection. The autofluorescence of the cells was eliminated by quantifying the signal from the unlabelled cells. For each sample 4-5 z-stacks were taken and 35-45 cells were quantified to study the cellular experiments.

### 3.5. In Vivo study

#### 3.5.1. Fish Strain

All experiments were done in *Tg(cldb::lyn-EGFP)* strain^37^, which marked the plasma membrane and helped in its visualization. For zebrafish maintenance and experimentation, guidelines recommended by the Committee for the Purpose of Control and Supervision of Experiments on Animals (CPCSEA), Govt. of India, were followed.

#### 3.5.2. DNA nanostructure treatment and Fixation

For the uptake of DNA nanocages, larvae at 72hpf were incubated in 300nM nanocage solution in E3 medium without methylene blue. To visualise the vesicles, Alexa Fluor 647 conjugated Wheat Gram Agglutinin – at the concentration of 5μg/ml was used as a tracer. The larvae were pulsed in this solution for 4 hrs followed by E3 (without methylene blue) washes and were then fixed in 4% PFA in PEMTT buffer (0.1M PIPES, 5mM EGTA, 2mM MgCl2.6H2O, 0.1% TritonX-100, 0.1% Tween 20, pH 6.8)^38^. Larvae pulsed with only tracer were used as control. The fixed embryos were kept overnight at 4 °C, followed by glycerol upgrade and storage at 4 °C.

#### 3.5.3. Image Acquisition and processing

Imaging was performed over the head (dorsal head periderm) of zebrafish larvae using Zeiss LSM 880 confocal microscope with an EC Plan 63x/1.40 oil immersion objective lens at 1.5x optical zoom with a Z-step of 0.37 μm. The images were taken at a resolution of 1024 × 1024. The pinhole values were kept at 1 Airy Unit. The PMT and laser power were kept constant while imaging the nanocages. ImageJ was used for image processing and analysis^39^.

### 4. Statistical Analysis

For in-vitro studies, the uptake of different DNA nanostructures in cells was quantified from the background subtracted cells using ImageJ software (NIH) and the image quantification using Graphpad Prism 8.0. For in vivo studies, the total number of WGA vesicles and the vesicles colocalising with the nanocages were counted manually and plotted. Experiments were not randomized. The investigators were not blinded to allocation during experiments and outcome. Mean values with associated standard deviation (SD) or standard error (SE) were used and are mentioned accordingly in the main manuscript. p values were calculated using one way ANOVA and two-tailed unpaired t-test on GraphPad Prism (confidence interval: 95%).

## 5. Acknowledgements

We sincerely thank all the members of DB group for critically reading the manuscript and their valuable feedback. AR, VM thank IITGN-MHRD, GoI PhD fellowship. PV acknowledges PhD fellowship from UGC-CSIR, India and IITGN for additional fellowship. DB thanks SERB, GoI for Ramanujan Fellowship, IITGN, for the startup grant, and DBT-EMR, Gujcost-DST, GSBTM and BRNS-BARC for research grants. MS acknowledges the support from TIFR-DAE (RTI4003;12P-121). Imaging facilities of CIF at IIT Gandhinagar are acknowledged.

## Supplementary Information

Sequence of all the oligonucleotides utilized to synthesise different DNA nanostructures.

**Table 1:**
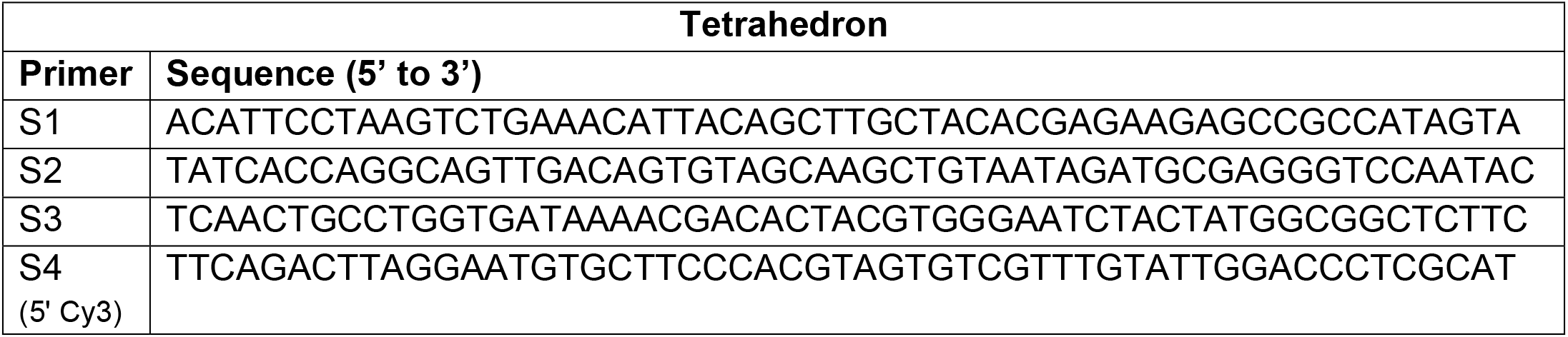
Tetrahedron.

**Table 2:**
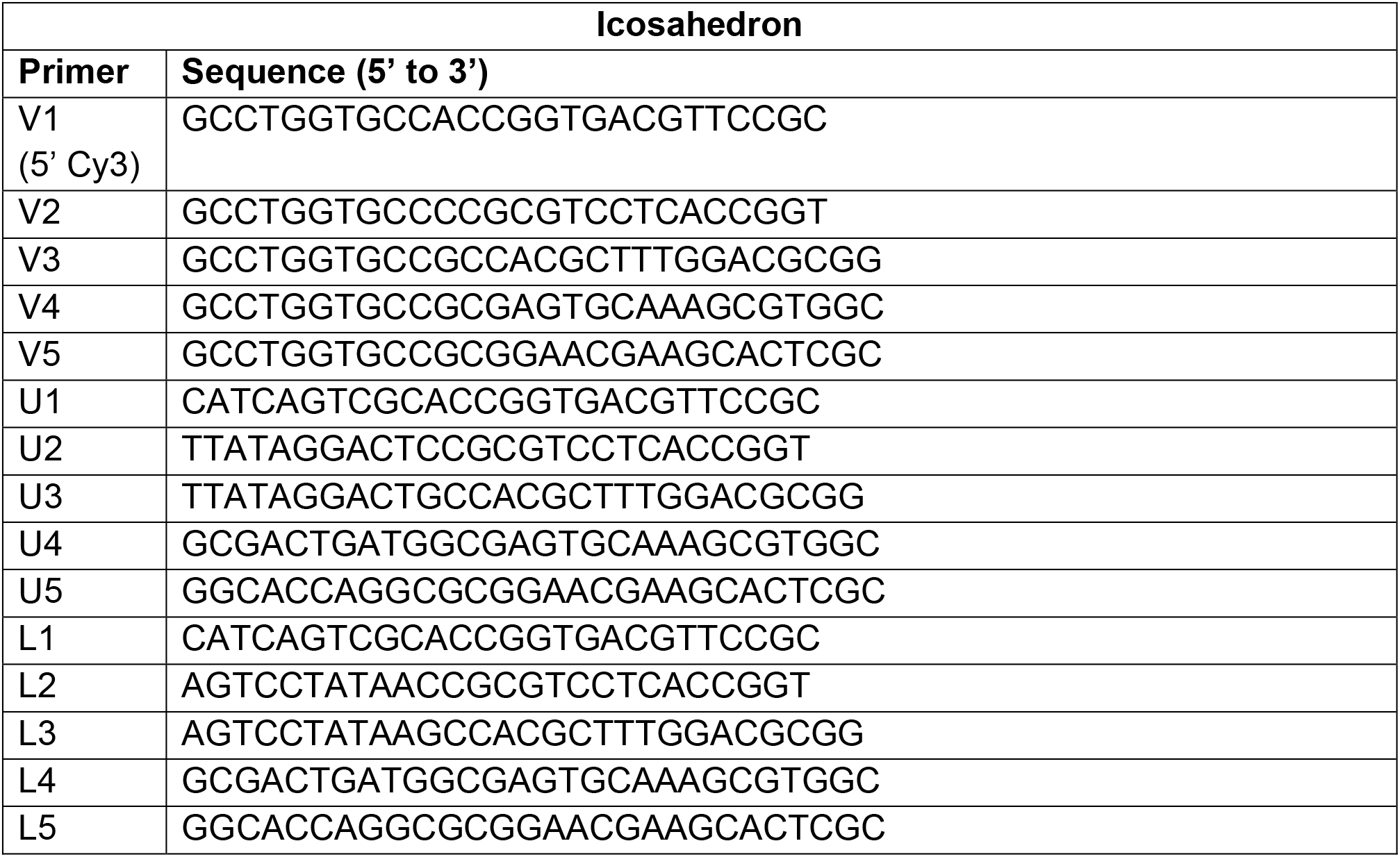
Icosahedron.

**Table 3:**
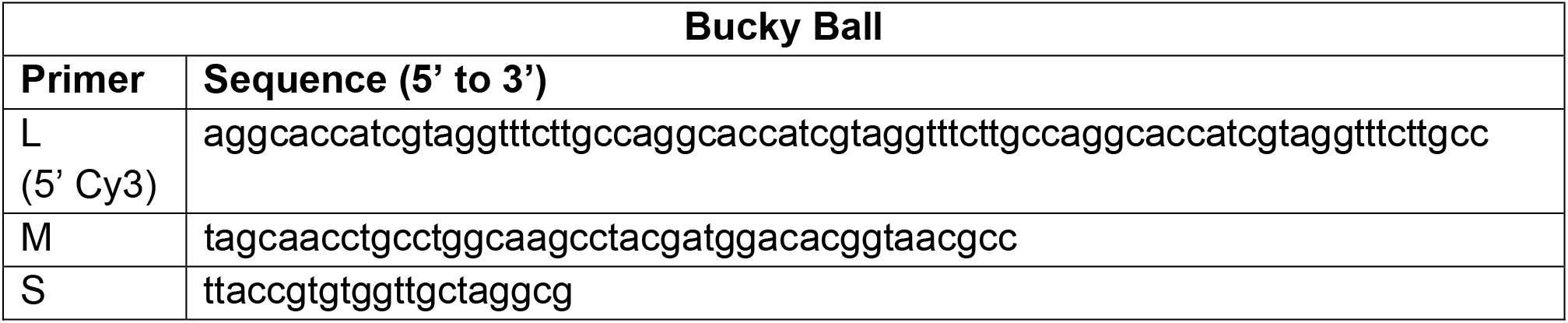
Bucky Ball.

**Table 4:** Cube.

| Cube |  |
| --- | --- |
| Primer | Sequence (5' to 3') |
| L<br>(5' Cy3) | Agg CAC CAT CgT Agg TTT TTCT TgC CAg gCA CCA TCg TAg GTT TTT<br>CTT gCC Agg CAC CAT CgT Agg TTT TTC TTg CC |
| MA | TAg CAA CCT gCC Tgg CAA gCC TAC gAT ggA CAC ggT AAC gAC -3' |
| SA | TTA CCg TgT ggT TgC TAg gCg |
| MB | TAg CAA CCT gCC Tgg CAA gCC TAC gAT ggA CAC ggT AAC gCC |
| SB | TTA CCg TgT ggT TgC TAg TCg |

**Figure S1:**
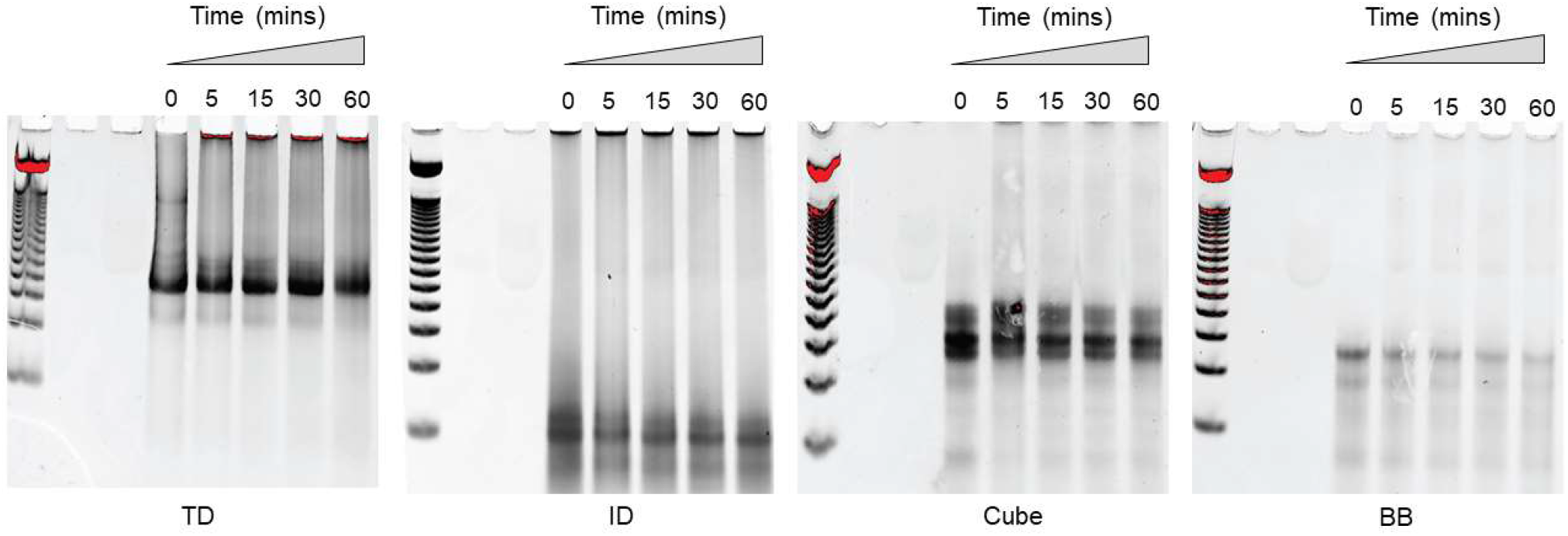
Stability of DNs in 10% FBS. DNs were incubated with non-heated FBS at 37°C for 5, 15, 30 and 60 mins. The integrity of DNs were examined by polyacrylamide gel electrophoresis. All DNs exhibited resistance to nuclease degradation for up to 30 mins.

**Figure S2:**
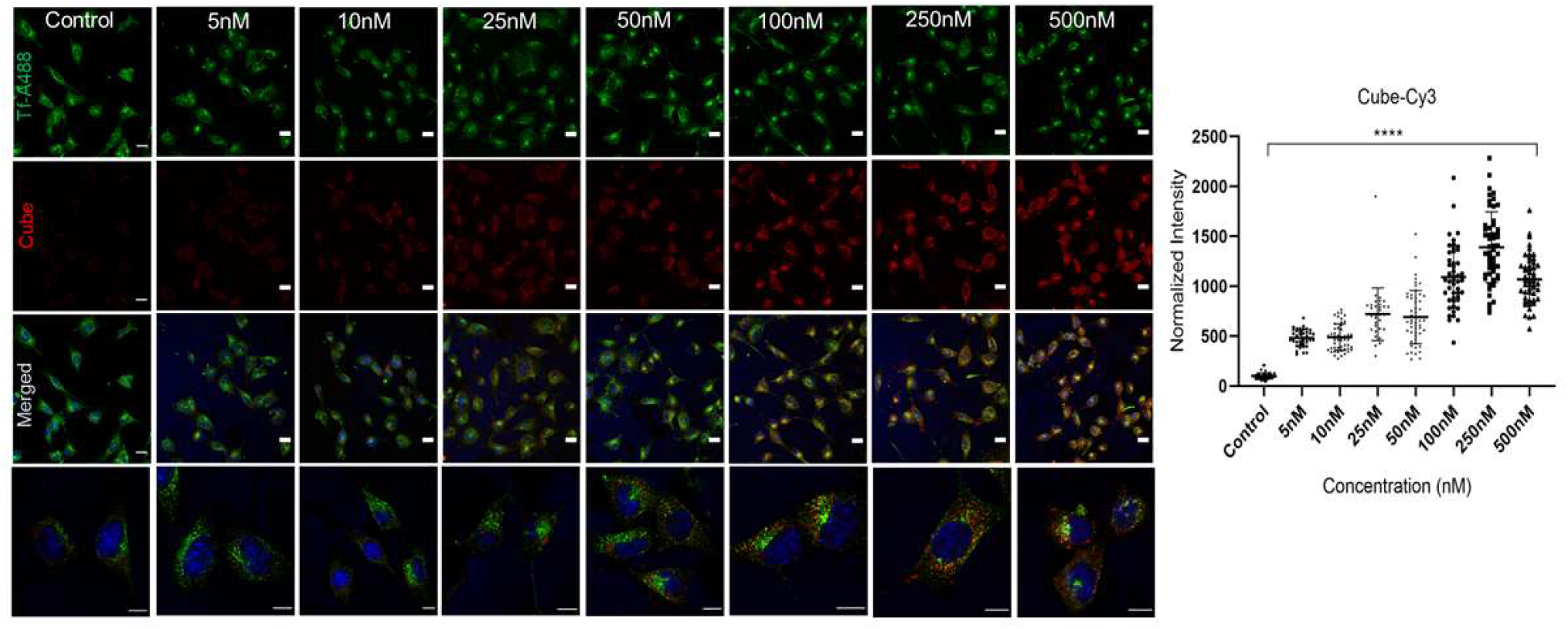
Concentration dependent uptake of DNA Cube. (a) Confocal images of MDA-MB231 cells treated with different concentration of Cy3 labelled DNA cube for 20mins at 37°C. Non-treated cells were considered as control. Green channel represents Tf-A488, red channel represents Cy3 labelled DNA cube ranging from 5nM to 500nM. Scale baer:10μm. (b) Quantitative uptake of Cube-Cy3 in MDA-MB231 cells from panel (a). Error bars represents Standard deviation. The normalized intensity was calculated from 40 cells (ordinary one-way ANOVA, P value < 0.0001****).

**Figure S3:**
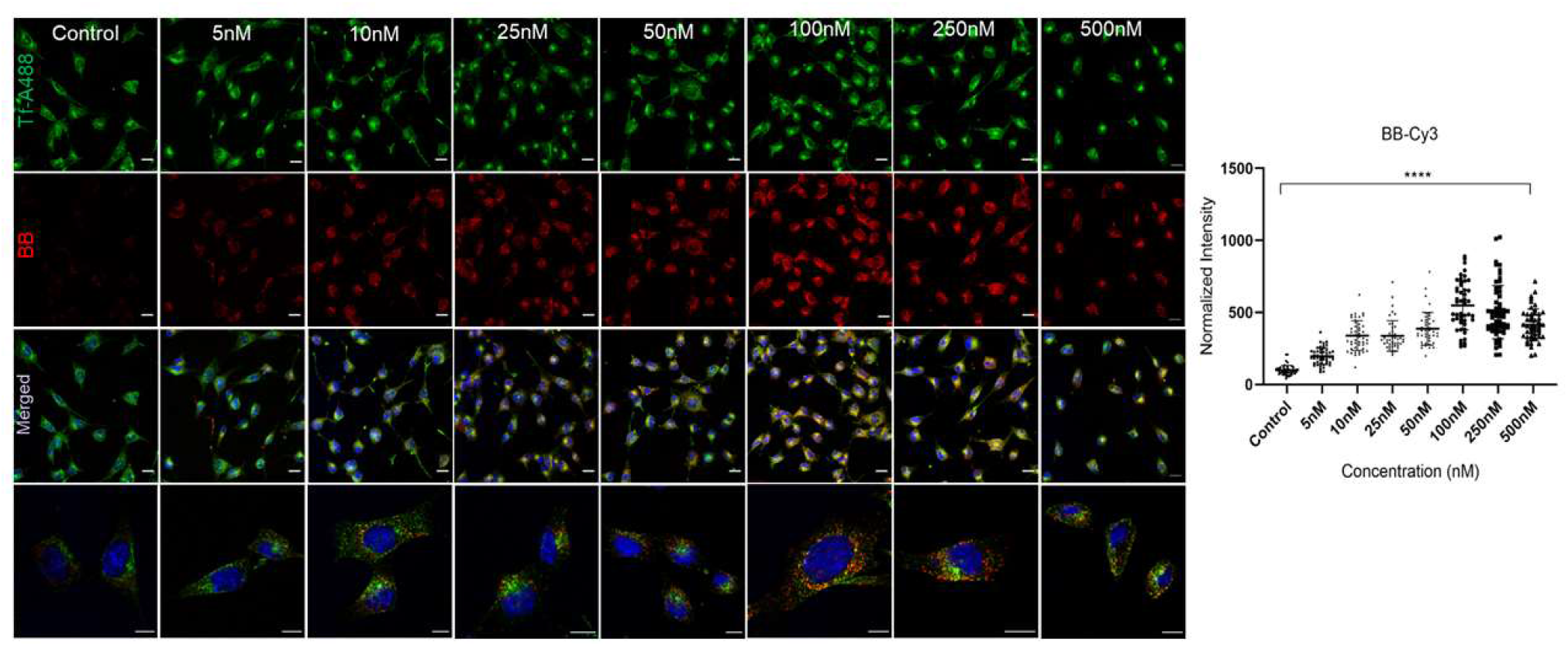
Concentration-dependent uptake of DNA BB. (a) Confocal images of MDA-MB231 cells treated with different concentrations of Cy3 labeled DNA BB for 20mins at 37°C. Non-treated cells were considered as controls. The Green channel represents Tf-A488, the red channel represents Cy3 labeled DNA BB ranging from 5nM to 500nM. Scale baer:10μm. (b) Quantitative uptake of BB-Cy3 in MDA-MB231 cells from panel (a). Error bars represent Standard deviation. The normalized intensity was calculated from 40 cells (ordinary one-way ANOVA, p-value < 0.0001****). Scale Bar: 10μm.

**Figure S4:**
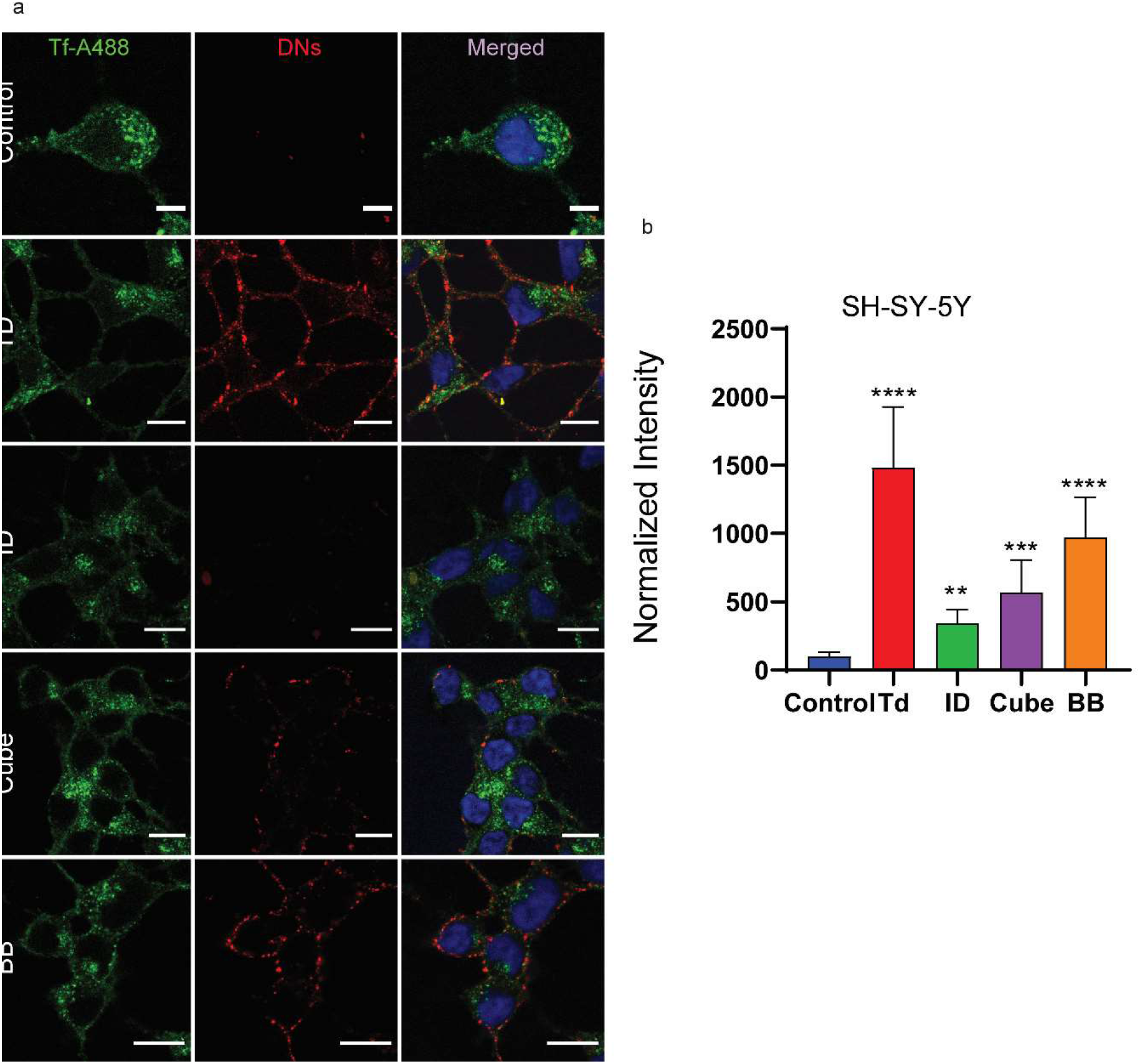
Cellular uptake of DNA nanostructures in SH-SY-5Y cells. (a) Confocal images showing the internalization of DNA nanostructures. Non-treated cells were considered as controls. Green colour indicated Tf-A488, red colour indicated Cy3 labeled DNA nanostructures. Quantitative analysis of internalized DNA nanostructures from cells in panel (a). The normalized intensity was calculated from 40 cells Error bars indicate the mean with s.d. (two-tailed unpaired t-test. **** p < 0.0001, *** p<0.0005, ** p<0.005). Scale Bar: 15μm.

**Figure S5:**
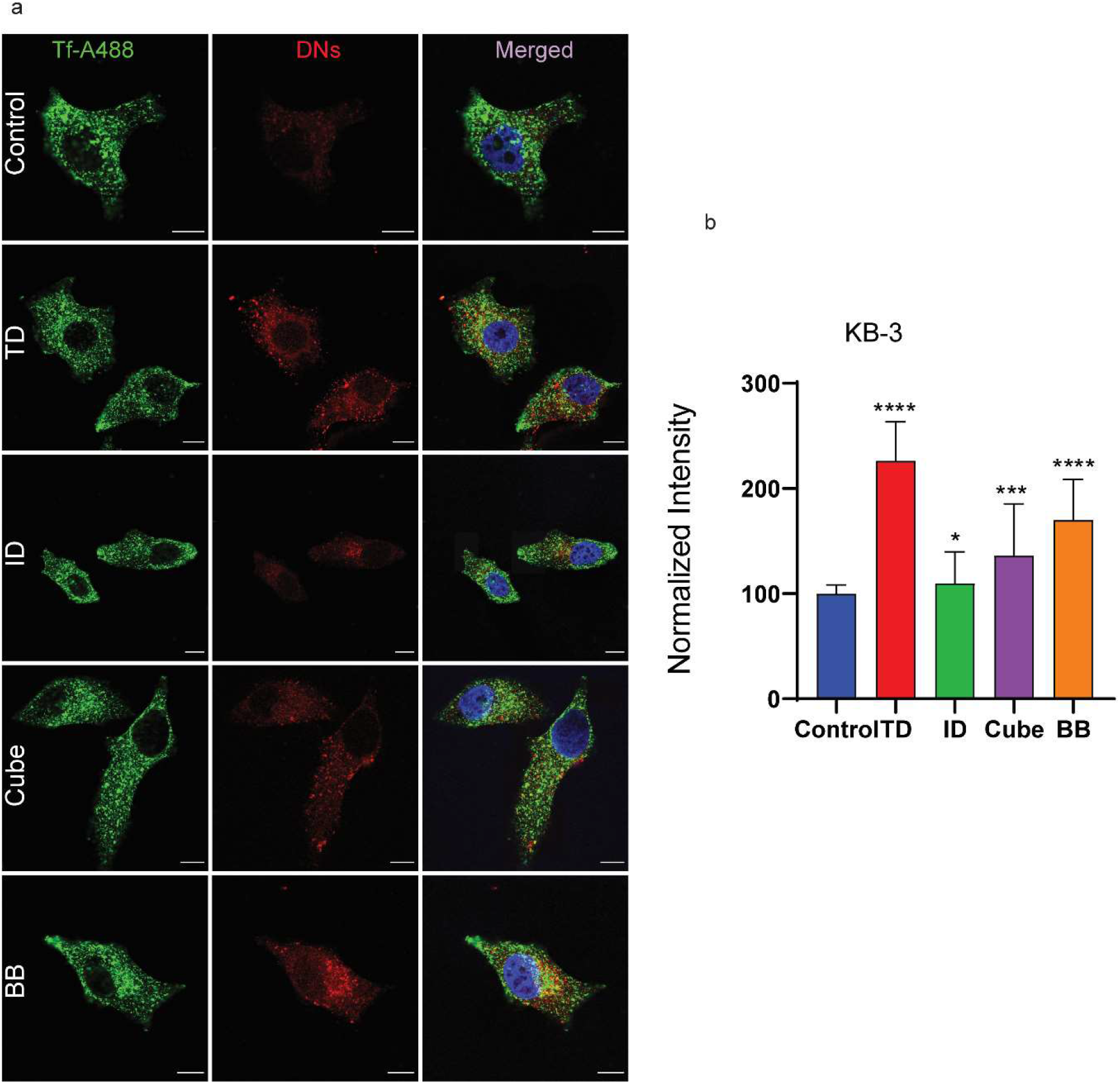
Cellular uptake of DNA nanostructures in KB-3 cells. (a) Confocal images showing the internalization of DNA nanostructures. Non-treated cells were considered as controls. Green colour indicated Tf-A488, red colour indicated Cy3 labeled DNA nanostructures. Quantitative analysis of internalized DNA nanostructures from cells in panel (a). The normalized intensity was calculated from 40 cells Error bars indicate the mean with s.d. (two-tailed unpaired t-test. **** p < 0.0001, *** p<0.0005, * p<0.0463). Scale Bar: 10μm.

**Figure S6:**
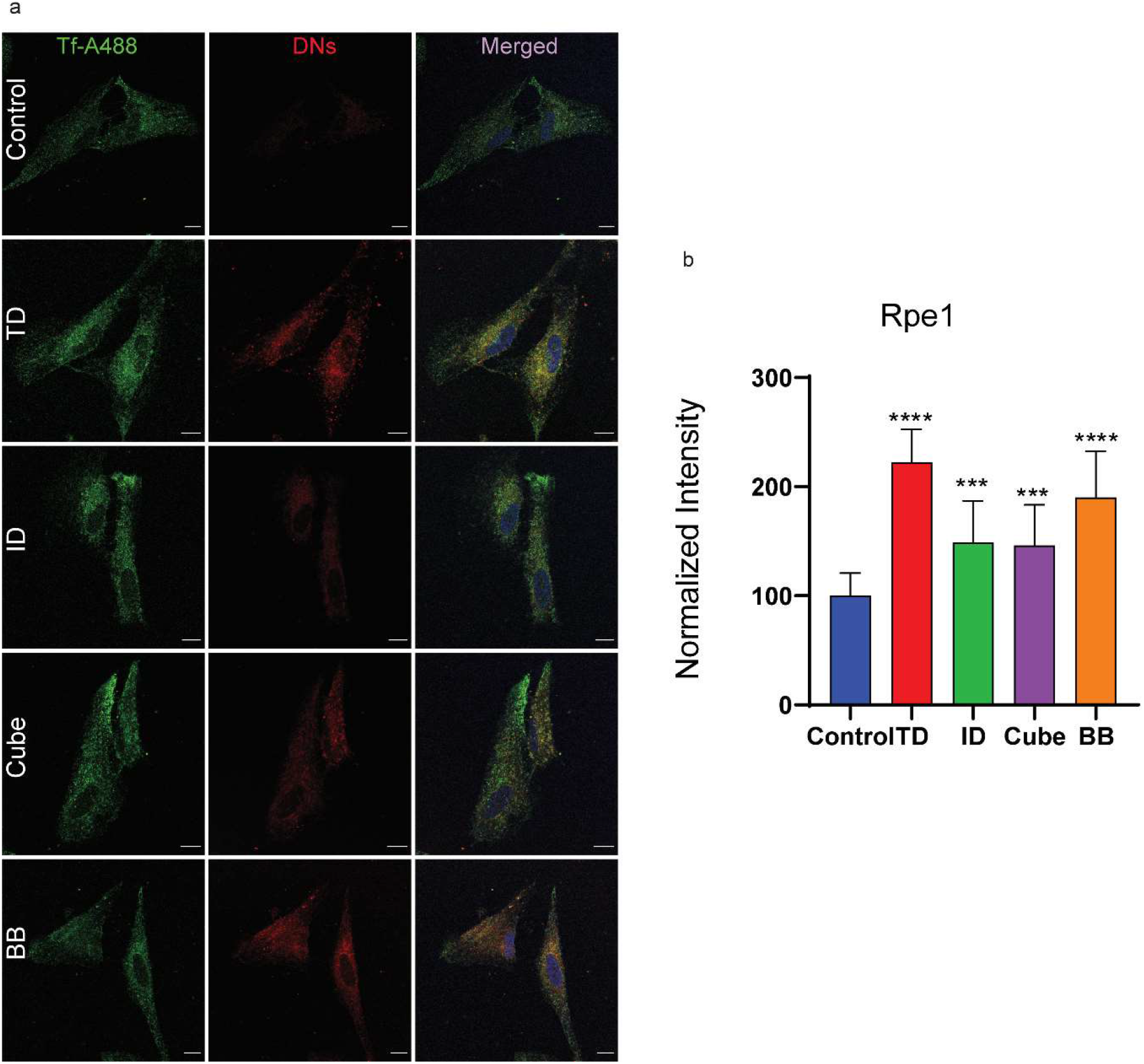
Cellular uptake of DNA nanostructures in Rpe1 cells. (a) Confocal images showing the internalization of DNA nanostructures. Non-treated cells were considered as controls. Green colour indicated Tf-A488, red colour indicated Cy3 labeled DNA nanostructures. Quantitative analysis of internalized DNA nanostructures from cells in panel (a). The normalized intensity was calculated from 40 cells Error bars indicate the mean with s.d. (two-tailed unpaired t-test. **** p < 0.0001, *** p<0.0005). Scale Bar: 10μm.

**Figure S7:**
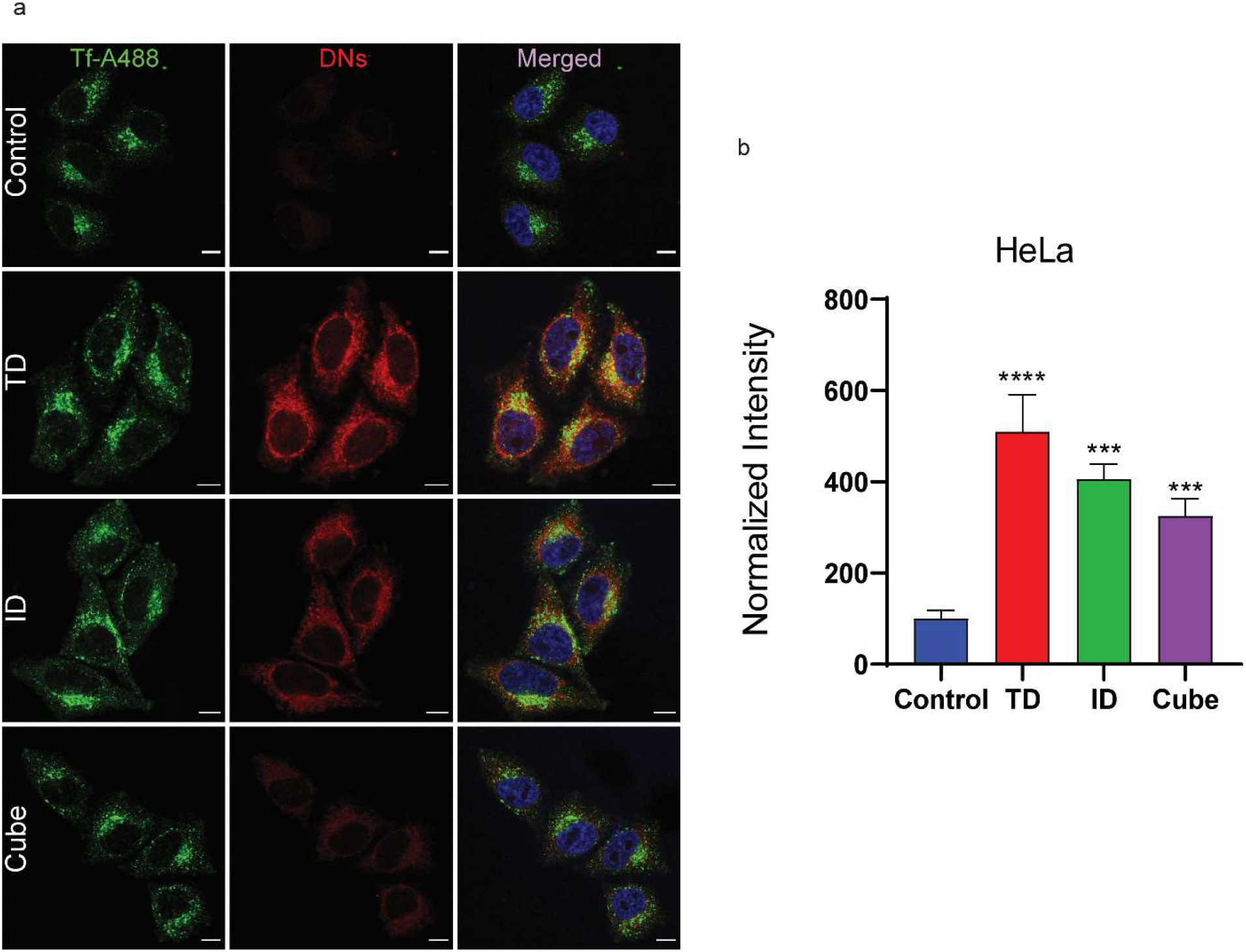
Cellular uptake of DNA nanostructures in HeLa cells. (a) Confocal images showing the internalization of DNA nanostructures. Non-treated cells were considered as controls. The green colour indicated Tf-A488, red colour indicated Cy3 labeled DNA nanostructures. Quantitative analysis of internalized DNA nanostructures from cells in panel (a). Error bars represent Standard deviation. The normalized intensity was calculated from 40 cells Error bars indicate the mean with s.d. (two-tailed unpaired t-test. **** p < 0.0001, *** p<0.0005). Scale Bar: 10μm.

**Figure S8:**
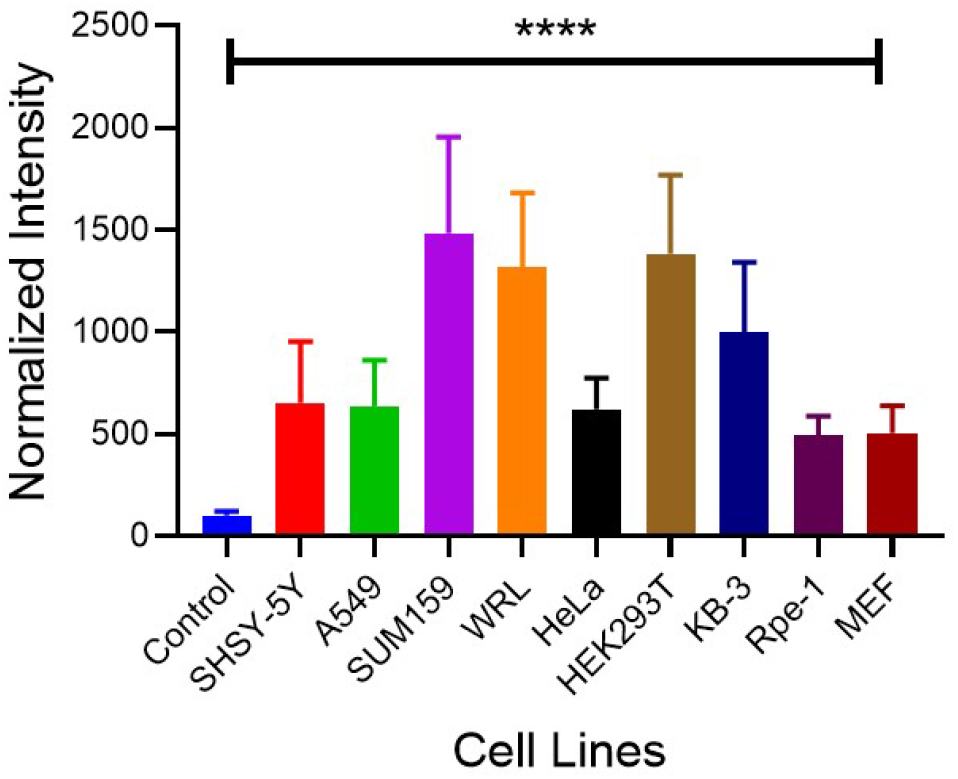
Cellular uptake of DNA TD in different cell lines. The internalization was maximum in (breast) SUM159 cells followed by (kidney) HEK293T, (liver) WRL, (oral) KB-3, (neuroblastoma) SH-SY-5Y, (lung) A549, (cervical) HeLa, (fibroblast) MEFs, and (retinal) Rpe1. The normalized intensity was calculated from 40 cells. Error bars indicate the mean with s.d. (two-tailed unpaired t-test. **** p < 0.0001.

**Figure S9:**
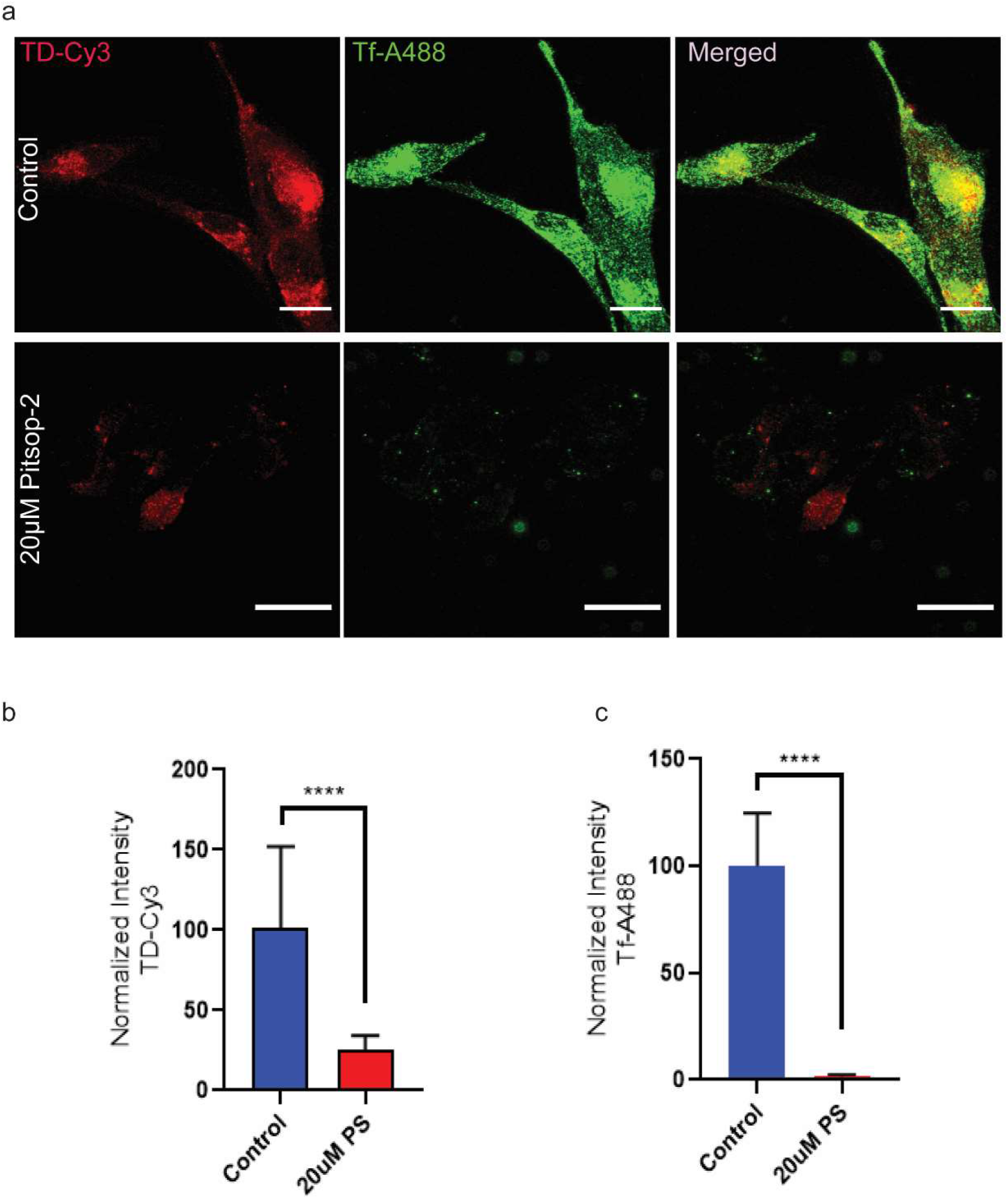
Cellular uptake of DNA TD via Clathrin mediated pathway in MDA-MB231 cells. Confocal images of MDA-MB231 cells showing the uptake of Tf-A488 in green channel, and TD-Cy3 in red channel for 20 min at 37°C in the presence or absence of 20 μM Pitstop-2. **(b)** Quantification of inhibition of CME pathway by Pitstop-2 for Tf-A488, and TD-Cy3 from cells in panel (a). Significant inhibition of TD and Tf upon treatment with pitstop-2. Error bars represent Standard deviation. The normalized intensity was calculated from 40 cells (ordinary one-way ANOVA, p-value < 0.0001****). Scale Bar: 20μm.

**Figure S10:**
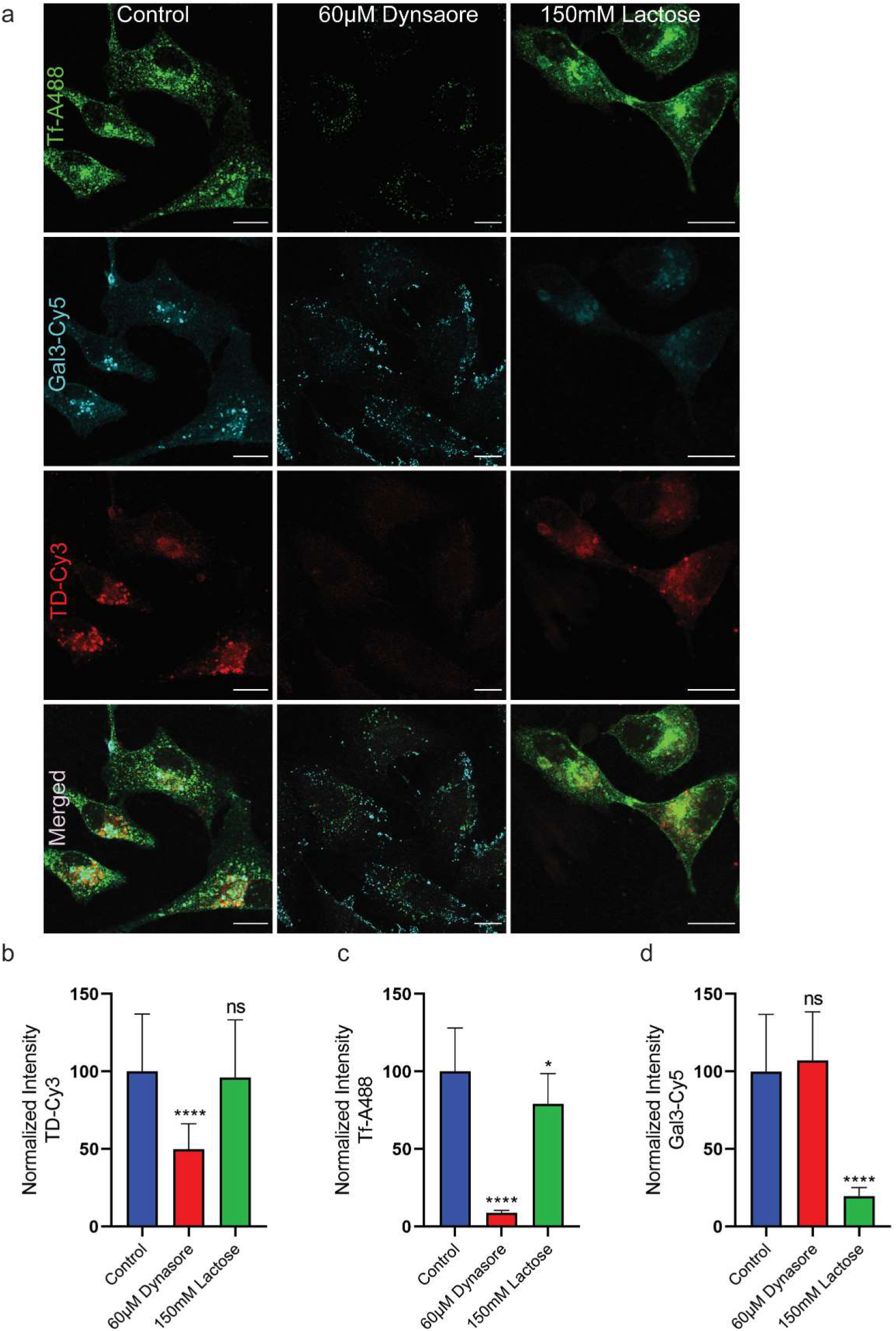
Validation of cellular uptake of DNA TD via clathrin mediated pathway using dynasore and lactose in MDA-MB231 cells. Confocal images of MDA-MB231cells showing the uptake of Tf-A488 in green channel, Gal3-Cy5 in cyan channel and TD-Cy3 in red channel for 20 min at 37°C in the presence or absence of 60 μM dynasore and 150mM Lactose. **(b)** Quantification of TD internalization by blocking the dynamin dependent pathway using dynasore and clathrin independent pathway using lactose. The inetrnalization of TD was significantly affected after dynasore treatment whereas no significant uptake in TD internalization upon treatment with lactose. **(c)** Quantification of Tf-A488 internalization which was used as an internal tracer of CME pathway upon treatment with dynasore and lactose. The internalization of Tf was significantly decreased upon dynasore treatment whereas the uptake was less affected in lactose treatment. **(d)** Quantification of Gal3-Cy5 which was used as a tracer for clathrin independent pathway. The internalization of gal3 was unaffected in case of dynasore and was significantly affected upon lactose treatment. Error bars represent Standard deviation. The normalized intensity was calculated from 30 cells (ordinary one-way ANOVA, p-value **** < 0.0001, *< 0.012). Scale Bar: 15μm.

**Figure S11:**
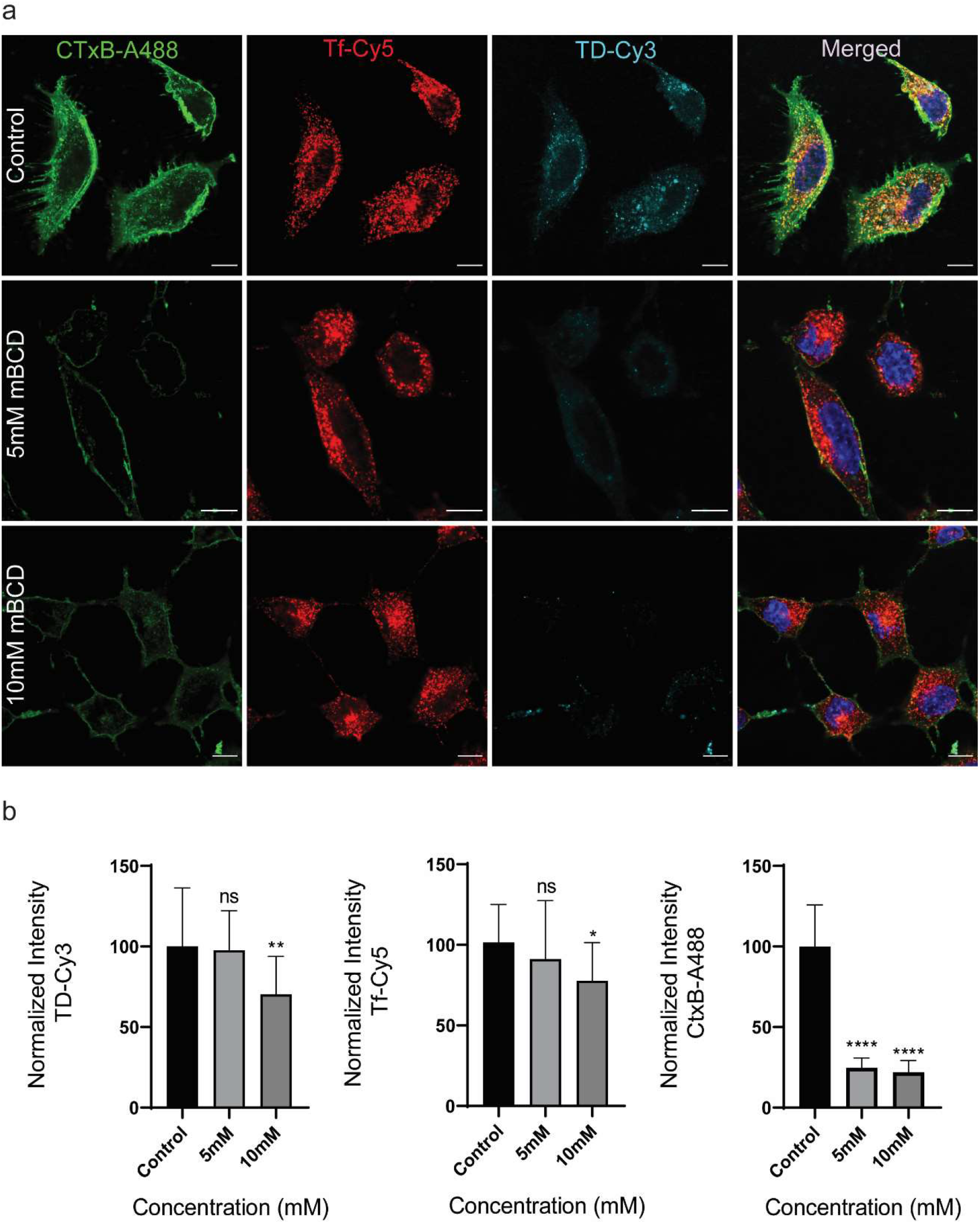
**(a)** Confocal images of MDA-MB231 cells showing the uptake of Tf-Cy5 in red channel, CTxB-A488 in Green channel, and TD-Cy3 in the cyan channel for 20 min at 37°C in the presence or absence of 5mM and 10mM mβCD. **(b)** Quantification of inhibition of caveolin pathway by mβCD for Tf-Cy5, TD-Cy3, and CTxB-A488 from cells in panel (a). Significant inhibition of CTxB-A488 at both concentrations indicates the caveolin pathway’s blocking but no significant inhibition of TD and TF at 5mM. The normalized intensity was calculated from 40 cells. Error bars indicate the mean with s.d. (two-tailed unpaired t-test. **** p < 0.0001, ** p<0.005, * p<0.05, ns non-significant >0.05).

